# Predictability of next elements in chimpanzee gesture sequences

**DOI:** 10.1101/2024.11.13.623362

**Authors:** Alexander Mielke, Gal Badihi, Ed Donnellan, Kirsty E. Graham, Chie Hashimoto, Joseph G. Mine, Alex K. Piel, Alexandra Safryghin, Katie E. Slocombe, Adrian Soldati, Fiona A. Stewart, Simon W. Townsend, Claudia Wilke, Klaus Zuberbühler, Chiara Zulberti, Catherine Hobaiter

## Abstract

Recent research has produced evidence for basic combinatorial abilities in the vocal systems of different animal species. Here, we investigate the structure of gesture sequences in Eastern chimpanzees (*Pan troglodytes schweinfurthii*) to detect whether gestural communication shows non-random combinations and how combinatorial rules influence predictability. Gesture, as compared to vocalization, offers greater flexibility in how signals are combined—for example overlapping in time — and as the parsing of signals into sequences is dependent on researcher decisions, we employ a multiverse approach, considering four different definitions of what constitutes a ‘sequence’ based on varying time thresholds. Our results indicate that sequences tend to be short (even with the most liberal time-window) and that transitions between some gesture types occur more frequently than expected by chance, with some transitions showing significant association across all time-windows. These transitions often involve repetition, suggesting persistence as a key aspect of chimpanzee gestural sequences. Information about previous gestures reduced uncertainty in predicting subsequent gestures. The order of gestures within sequences appears to be less critical than their cooccurrence, challenging assumptions based on the linear patterning derived from vocal communication. Our findings highlight the importance of methodological choices in sequence definition and suggest that chimpanzee gestural communication is characterised by a mix of predictability and flexibility, with implications for understanding the evolution of complex communication systems.

## Introduction

Despite substantial differences in the expression of human language and other animals’ communication systems, research has increasingly uncovered similarities in structure (e.g., Berthet et al., 2023), providing evidence that there are fewer qualitative differences than previously envisioned. Animal communication systems are characterised by a rich array of structural properties, including evidence of hierarchical organisation (ten Cate & Okanoya, 2012; Weiss et al., 2014), non-Markovian dependencies (Kershenbaum et al., 2014), and combinatoriality (Coye et al., 2018; Engesser et al., 2016; Oña et al., 2019; Walsh et al., 2023). Combinatoriality, the ability to predictably produce more than one element either in sequence or simultaneously, has the potential to dramatically increase the amount of information that can be transmitted with a finite repertoire, especially when combinations can take on meaning that is distinct from the constituting parts (compositionality). Many studies of combinatoriality in animal communication have been conducted on systems that are not considered to have meaning-bearing signal elements (in the linguistic sense), for example the song-like sequences of whales, birds, and some primates (Allen et al., 2019; Berwick et al., 2011; Clarke et al., 2006). Research on meaning-bearing signals has often focussed on restricted sets of highly specific meaning-bearing systems, such as alarm calls (Coye et al., 2018; Engesser et al., 2016; Suzuki et al., 2019), usually favouring vocal communication. Using specific contexts to test animals’ ability to combine signals non-randomly has the advantage of reducing noise, but it gives us little indication of how these skills (if they exist) influence the communication system as a whole. Here, we use the largest database of coded chimpanzee gestures to test whether there are non-random transitions between gesture actions (n-grams), based on one or two previous elements in a sequence, and how any combinatorial rules underlying these transitions influence the predictability of sequences.

There is ample evidence in singing species that individuals combine signals into predictable structures, the acquisition of which often involves evidence of individual learning and culture (Allen et al., 2019; Berwick et al., 2011; Clarke et al., 2006; Garland et al., 2011; Kershenbaum et al., 2012). Datasets for this approach can be very large, as song sequences are typically longer than those in other forms of non-human communication, so probabilistic transition patterns can be established using methods imported from natural language processing (Kershenbaum et al., 2016). For meaning-bearing communication, we have evidence from birds’ and primates’ vocalisations using observational and experimental approaches that identify predictable transitions between elements. These studies often focus on identifying specific combinations that can then be tested experimentally (e.g., Dutour et al., 2019; Schlenker et al., 2016; Spiess et al., 2022; Suzuki et al., 2016). In non-vocal communication, datasets have been more limited—there is at least preliminary evidence for predictable sequences or co-occurrences in gestural communication (Genty & Byrne, 2010), facial signals (Oña et al., 2019), and multimodal signals (Aychet et al., 2021; Muschinski et al., 2023; Wilke et al., 2017), but a particular challenge in these systems is that the repertoires of units are often far larger than found in vocal communication (Hobaiter & Byrne, 2011b) and the ratio of distinct communication units to sample size has made it difficult to understand the system as a whole, rather than for small subsamples of common units.

Here, we analyse the structure of chimpanzee gesture sequences as a system, rather than focusing on a single context, using an uncommonly large dataset (Grund et al., 2023; Mielke et al., 2024). Given their evolutionary proximity to our own species, their large repertoire of gestural units, the diversity of contexts in which they communicate, and existing research on the meaning, flexibility, and intentionality of this system, chimpanzees are a prime target for this type of study. Chimpanzee gestural communication is frequently sequential by nature, partly due to chimpanzees often persisting and elaborating in order to meet their goals (Hobaiter & Byrne, 2011b; Leavens et al., 2005; Rodrigues & Fröhlich, 2021). However, sequences are often short, with a previous study finding that almost 80% of ‘rapid-fire sequences’ (those made with less than a second between units) were composed of only 2-elements (Hobaiter & Byrne, 2011a). In rapid-fire sequences, repetition of the same gesture is relatively rare, but it becomes increasingly common when longer pauses between gestures, including periods of response waiting, are considered (Hobaiter & Byrne, 2011a). Response waiting is a characteristic of intentional imperative communication in which the signaller seeks to achieve a particular goal with respect to a specific recipient (Bates et al., 1975) — as a result, as opposed to continuing to broadcast information, the signaller emits a signal and then waits to see whether the recipient will respond, in order to determine whether they should produce additional signals. Younger individuals use more and longer sequences, possibly because these are more common in play, but also potentially to explore their large gestural repertoires (Hobaiter & Byrne, 2011b). There is divergent evidence about the impact of gesture sequences on communicative efficiency, with some studies failing to identify a clear impact (Genty & Byrne, 2010) while others suggesting that stringing gestures together increases success, but only because it increased the likelihood of incorporating an inherently more successful gesture unit (Hobaiter & Byrne, 2011a). Transitions between gesture units have been reported in lowland gorillas, showing a number of regular transitions, many of which were repetitions (Genty & Byrne, 2010). Gesture actions in lowland gorillas form clusters, with physical contact gestures within play often occurring in an unordered cluster and play initiation/regulation gestures occurring in an ordered cluster (Genty & Byrne, 2010), a pattern also seen in chimpanzee play (Mielke & Carvalho, 2022). Combinations of facial signals with gestures can alter their meaning in predictable ways, with bared-teeth faces impacting whether a gesture elicits an affiliative response (Oña et al., 2019).

More evidence of predictable rules in chimpanzee sequential signalling derives from vocal communication. Sequential utterances are present from birth (Soldati et al., 2022), and sequence production increases in ‘complexity’ (in terms of their length and diversity) throughout infancy, coinciding with a diversification of social interactions (Bortolato et al., 2023). Evidence from different populations now shows that call combinations remain common in adulthood (Crockford & Boesch, 2005; Leroux et al., 2022). Here analyses have focused on how sequences change the information encoded in both specific contexts (e.g., food detection, Leroux et al., 2021) and the repertoire more widely (Girard-Buttoz et al., 2022; Leroux et al., 2022). These datasets show that chimpanzees can combine units flexibly (i.e., each unit occurs in sequence with multiple others), that there are production biases for unit order (AB occurring at different rates than BA), and evidence of the recombination of two-unit combinations (bigrams) in three-unit combinations and larger structures (Girard-Buttoz et al., 2022; Leroux et al., 2021). However, in many vocal studies, vocalisations are lumped into small repertoire sets of higher-order call types. Recent work provides ample evidence that calls are graded and— importantly—encode specific context at a much finer-grained scale (Crockford et al., 2018; Slocombe & Zuberbühler, 2005). As a result, it remains challenging to assess the true flexibility of the system: it might be that ‘hoos’ in general (or ‘screams’ or ‘barks’) are combined with a number of other units, but more specific units such as ‘travel hoos’ only occur in fixed combinations, belying the appearance of recombination.

Adding to the challenge of studying combinations, in the broader study of non-human communication, there is no natural definition of what constitutes a cohesive sequence of units. This ambiguity has led to a proliferation of different cut-off values to differentiate between one sequence and another—in the case of analyses of bird or whale songs, for example (Allen et al., 2019), sequences are anything that happens between the start and end of the song. In contrast, gesture researchers have historically applied different thresholds; for example, considering units to be in the same rapid-fire sequence when the following gesture occurs within one second of the end of the previous one (Hobaiter & Byrne, 2011a), a rule that has also been adopted in some vocal research (Girard-Buttoz et al., 2022), while other researchers have advocated a data-driven threshold detection approach (Aychet et al., 2021). Rapid-fire sequences in previous studies were differentiated from more extended sequences (sometimes called “bouts”) that represent the addition of further units for the purpose of persistence or elaboration following the failure of an initial communication (Leavens et al., 2005). The distinction between sequences and bouts was inspired by the idea of response waiting. If response waiting does define a different *type* of sequence, the different time-windows might contain different structural properties; for example, rapid-fire sequences (those without response waiting between gestures) might have distinct combinatorial rules as compared to sequences that include response waiting between signals (Hobaiter & Byrne, 2011a).

Importantly, while strictly vocal signals must almost always occur as separate signals produced in a linear sequence, gestures and multimodal sequences can be produced simultaneously using multiple limbs. This feature offers intriguing possibilities for encoding structure in dimensions such as space, as well as in time, as seen in signed languages (Armstrong et al., 1994; Sandler & Lillo-Martin, 2006; Stokoe, 1980) and human facial signals (Trujillo & Holler, 2024). Thus, we can define ‘sequences’ in a gestural context following different rules that might have both biological implications (because individuals communicate different pieces of information at different speeds, including units that overlap in time) and statistical implications (because more liberal sequence definitions will create longer sequences and therefore increase sample size). To address this, we take a multiverse approach (Steegen et al., 2016), conducting the same analytical steps using datasets following different preprocessing decisions to reduce researcher biases. For all analyses addressing first order transitions (A leading to B), we will produce results based on 4 different definitions of ‘sequence’ and use differences in outcomes to understand potential reasons for the differences in time thresholds and/or production speed. For analyses addressing higher-order transitions (AB leading to C), we will rely on the most liberal definition of sequence (here any gesture that occurs within 5 seconds of the end of the minimal action unit of the previous gesture), which adds noise but at the same time substantially increases the sample size for sequences above the length of 2 elements.

We further the study of sequence use in non-human species by analysing the largest dataset of chimpanzee gestures. We ask three questions: 1) **Are there non-random transitions between individual units and higher-order combinations?** We test this question by comparing observed transition probabilities between antecedent and consequent gestures with expected probabilities based on a null model of random distribution of elements (Aychet et al., 2021; Bosshard et al., 2022; Mielke & Carvalho, 2022). Thus, we establish which gesture actions are connected to each other, whether individual gesture actions have deterministic connections to specific subsequent gestures, and whether we find highly structured networks of gestures that occur together. We hypothesise that, as in vocalisations, there are bigrams and trigrams of gesture actions that occur next to each other in sequences more than would be expected if gesture actions were randomly distributed within or between sequences [P(B|A)] (Bosshard et al., 2022; Girard-Buttoz et al., 2022; Leroux et al., 2022). We hypothesise that with an increasing number of previous units, the number of significant n-grams decreases, but that we still find higher-order significant collocations [P(C|AB)] (Girard-Buttoz et al., 2022). We predict that, when analysing the transition network, we can identify clusters of gesture action that occur together frequently, even if their exact temporal order might not be relevant (Mielke & Carvalho, 2022). We also specifically compare the solitary gesture network and rapid-fire sequence network, as these potentially represent two distinct systems of combinatoriality with separate use: solitary gestures here are defined by individual elements separated by clear response waiting, so the sequence is clearly influenced by partner responses. Rapid-fire sequences are so fast that signaller planning should precede the partner response. **2) Does sequence order matter?** Order effects are important for establishing combinatoriality because they rule out a simpler alternative - that sequences are strings of a number of elements with the same meaning that are randomly put together until the partner reacts appropriately. We predict that we observe two-element pairs where the conditional probability of A following B exceeds what would be expected if sequences were randomly strung together. **3) How do these rules influence predictability?** We test this using entropy and classifier accuracy to establish the level of predictability and order in the system. Both provide us with estimates of the predictability of the system as a whole. For all models of Markov transitions (A leading to B), we employ four time-windows (below). We predict that knowing the previous unit(s) in a sequence improves the predictability and order in the system (quantified using entropy and classifier accuracy). We predict that order matters— randomising the order of A and B in the latter example should reduce predictability. When specifically focusing on cases where we predict a unit based on two preceding units, we predict that knowing both units [P(C|AB)] will improve prediction accuracy over just knowing either of them [P(C|A) or P(C|B)].

## Methods

### Data

The dataset contains 7,749 gestures from 5 communities of East African chimpanzees (*Pan troglodytes schweinfurthii)* (Mielke et al., 2024), structured in varying numbers of sequences of between one and 15 units, depending on the definition (Table 1). The coding scheme has been described in detail (Grund et al., 2023). The 92 gesture actions (distinct and delineated types of gestures) used in the original coding scheme were lumped based on predefined rules (see Mielke et al., 2024) to reduce the occurrence of rare cases, with gesture actions that occurred fewer than 10 times excluded (Mielke et al., 2024). We retained 42 gesture actions in the current analyses. Each gesture action has a clearly defined Minimum Action Unit (MAU) based on the minimum information necessary to distinguish between gesture actions, starting from the moment the individual moves the body part and finishing when the gesture action is fully in place (Grund et al., 2023). Note that gesture actions can continue to be held in place or repeated (through the use of an optional hold or repetition phase, *sensu* Kendon, 2004) beyond the end of the MAU, followed by recovery of the articulator to rest. Sequences were defined based on 4 distinct time-windows (Fig. 1), going from the broadest to the most restrictive: a) gestures that occurred within 5 seconds after the end of the MAU of the previous unit (*‘5 seconds*’); b) gestures that occurred within 1 second after the end of the MAU of the previous unit (previously termed ‘*rapid-fire sequences*’); c) gestures that occurred with overlap of their MAU (‘*overlap*’); d) gestures that were separated by at least one second, but occurred within the same 5 second windows (*solitary-gestures-plus-waiting*’). Category d) represents sequences where the choice to add a gesture to the sequence can be most clearly related to the recipient’s actions, while a) through c) at least partially contain sequences where individuals choose gestures in advance. Repetitions of the same gesture were treated depending on structural features (Grund et al., 2023): rhythmic repetition of the same gesture action (i.e. the same action repeated at the same pace in a continuous movement) was treated as a single occurrence with modifying information, while non-rhythmic repetition of the same gesture action (i.e. the action is repeated but the pace of motion differs and/or is not continuous) was treated as two distinct occurrences. This distinction differs from preprocessing applied to vocal sequences, where any repetition was treated as the same unit (Girard-Buttoz et al., 2022).

**Figure 1:**
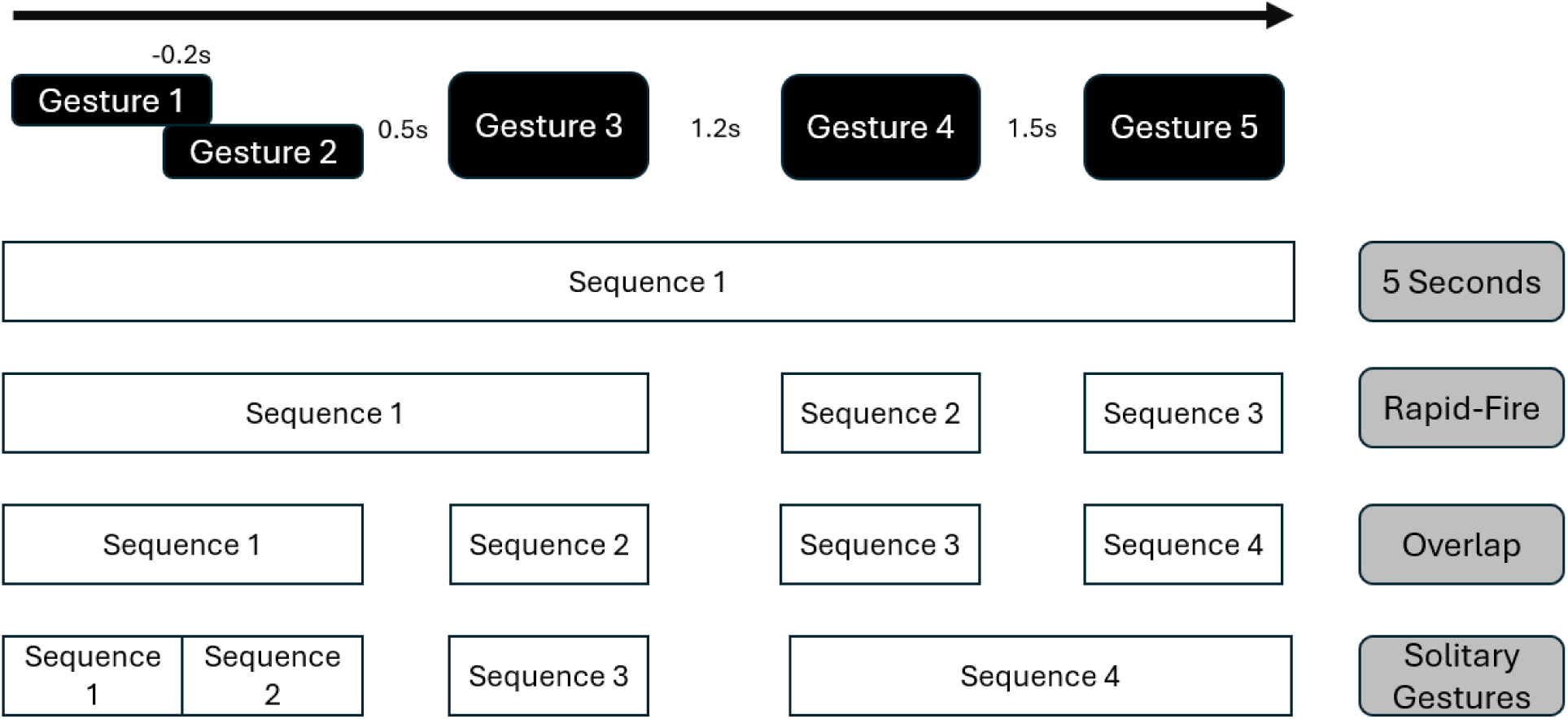
Difference between time-windows in classifying sequences: the 5 seconds time-window classes all five gestures as part of the same sequence based on the time between elements. The rapid-fire time-window classes gestures starting within 1 second of the previous MOU end as part of the same sequence. The overlap time-window only classes those with temporal overlap as part of the same sequence. The solitary-gesture-plus-waiting time-window classifies those that are the only gesture to occur in a 1 second window but are less than 5 seconds apart as belonging to the same sequence.

**Table 1:**
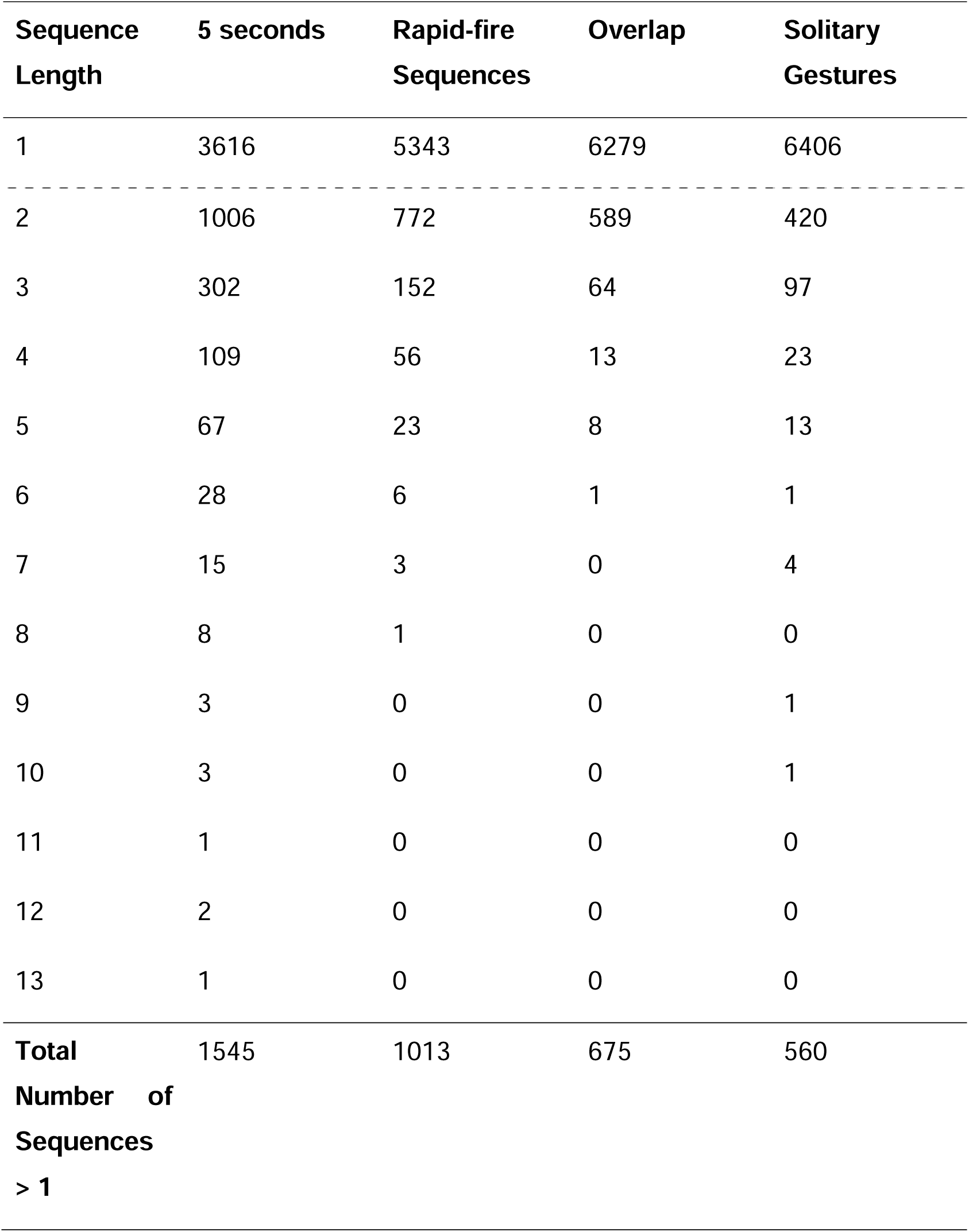
Number of sequences with a given number of units depending on time-window.

For next unit prediction, we broke longer sequences into n-grams of consecutive units. As this constitutes a very small dataset for the higher-order n-grams (e.g., 2,586 bigrams; 1,041 trigrams; and 502 4-grams for the 5-seconds sequences), we used bootstraps of the conditional probabilities to check whether analyses of trigrams and 4-grams could provide any information at all. We sampled sequences with replacement and plotted the range of conditional probabilities for each n-gram (Fig. 2). Because the 4-grams were unstable (most probabilities could take any value between 0 and 1), we dropped them from further analyses, focusing on bigrams and trigrams and acknowledging that the latter show considerable sample error. For the other time-windows, higher-order n-grams were too rare, so we only reported bigrams for these.

**Figure 2:**
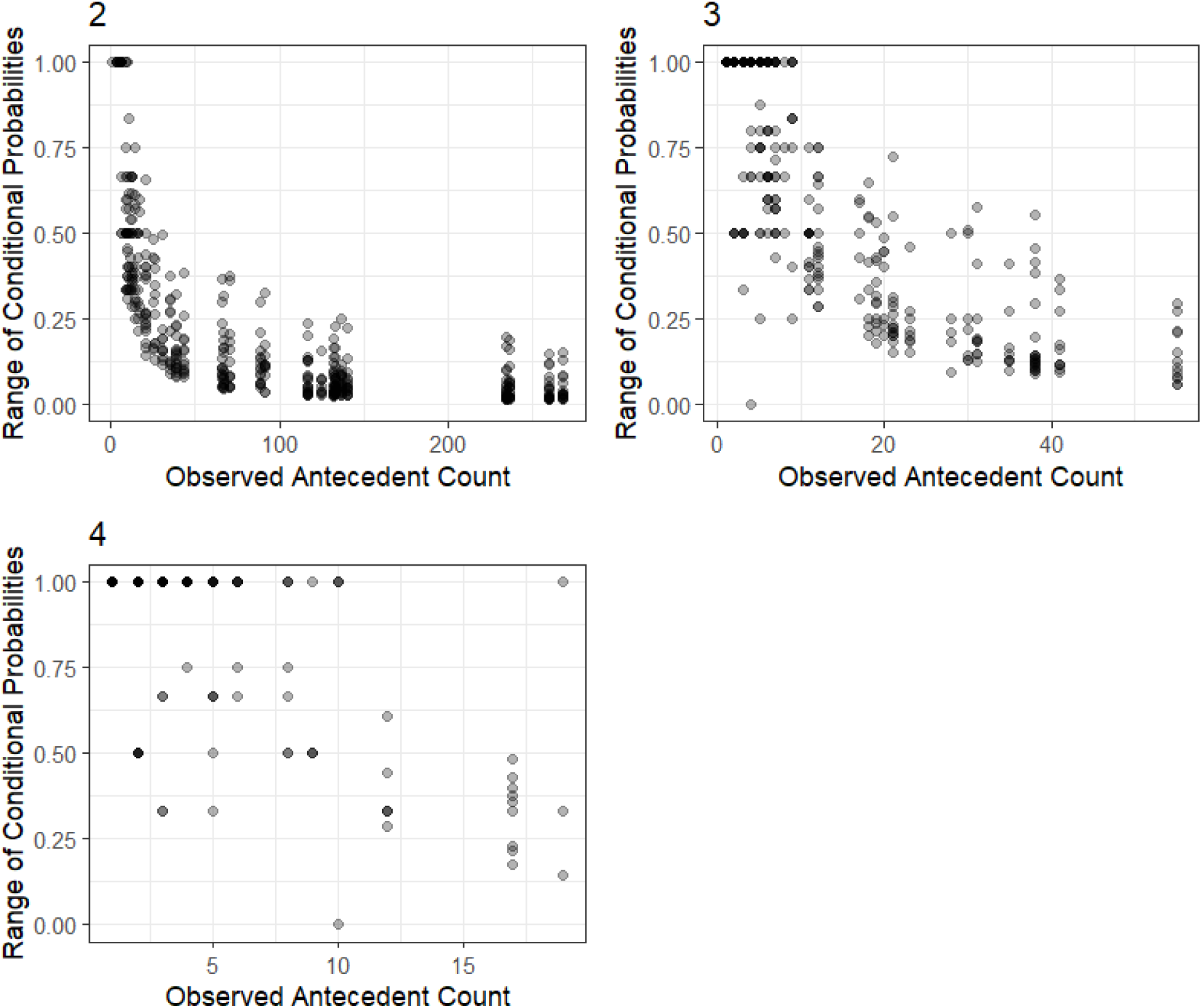
Variability of conditional transition probabilities based on the occurrence frequency of the antecedent unit for bigrams (one antecedent), trigrams (two antecedents), and 4-grams (three antecedents). The y-axis depicts the difference between the maximum and minimum conditional probability for that transition across 1,000 bootstraps.

All analyses were conducted in R and R studio (R Development Core Team & R Core Team, 2020) using the ‘tidyverse’ (Wickham et al., 2019) and ‘tidymodels’ (Kuhn & Wickham, 2020) environments.

### Significant transitions

To test which transitions occur at higher probabilities than expected, we calculated the conditional probabilities of a consequent unit given one (P(B|A)) and two (P(C|AB)) antecedents, corresponding to bigrams and trigrams. To calculate which transition probabilities occurred at significantly higher probabilities than expected, we compared the observed values to a null model generated by shuffling gesture actions across sequences while controlling for the number of units per sequence. ‘Significant’ transitions in this case are those that have a higher conditional probability than would be expected if units were randomly distributed (observed transition probability > 950 out of 1000 permutations) while accounting for the base probability of each gesture type. We only report transitions that occurred at least five times in the dataset, as conditional probabilities were nearly random for less frequent transitions (Fig. 2). Because we calculate these transitions across four different time-windows, we report and plot transition patterns that are significant and occur at least 5 times in at least three out of the four datasets, with the rationale that these will be the transition patterns that are most robust to researcher decisions. We also specifically compared the solitary-gesture-with waiting network and rapid-fire network, as these potentially represent two distinct systems of combinatoriality with separate use: solitary gestures here are defined by individual elements separated by clear response waiting, so the sequence is clearly influenced by partner responses. Rapid-fire sequences are so fast that signaller planning should precede the partner response. To find higher-order structure of transition patterns, we created the bigram transition network, representing each gesture action as a node and each significant bigram transition that occurred at least 5 times as an edge using igraph in R (Csardi & Nepusz, 2006). Clusters of elements were determined using the ‘cluster_optimal’ community detection algorithm (Brandes et al., 2008). Loops (conditional probabilities between gesture actions and themselves) are meaningful and relevant here, so we present them graphically.

To test whether there are dyads of units with a prescribed order (AB rather than BA), we randomised gesture action within sequences while keeping the identity of actions within sequences the same. ‘Significant’ transitions in this case are those that are observed more often than would be expected if order within sequences did not matter (transition probability > 950 out of 1000 permutations).

### Entropy and prediction accuracy

We use two measures to quantify how ordered next-element prediction is in this system. These provide similar information about how easy it would be for an individual to predict the next gesture in a sequence without additional information other than the previous gestures, but we promote diversity of methods to rule out that results are based on researcher choices.

a. Conditional entropies of transition probabilities: across time-windows, we calculated the observed conditional entropy for the bigrams. For the 5-second sequence, we also calculated the conditional entropy for the trigrams, plotting the entropy development (McCowan et al., 1999). Entropy was calculated using the ‘infotheo’ package in R (Meyer, 2022) and reported in bits. The entropy was calculated as:

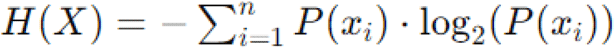

Lower entropy indicates less uncertainty and more order. To account for the different sample sizes at different levels and time-windows, we compared the observed entropies with those expected if gesture actions were randomly distributed across sequences by calculating the entropy ratio (observed entropy/random entropy), with values closer to 0 indicating that, knowing previous actions leads to higher predictability and values closer to 1 indicating random distribution. Because some time-windows have much smaller datasets, we show the development of the entropy ratio for randomly selected subsamples of each time-window (Mielke et al., 2021). To account for the possibility that previous units might reduce uncertainty but that the unit order might not matter in a trigram, we also added ‘alphabetised order’ for AB-C: treating AB and BA the same, we reduce the overall number of possible transitions; if we find lower entropy for this, by level, than for the ordered transitions, we would assume that order does not matter and adds noise. We calculated entropies for 100 bootstraps (sampling with replacement) to give a sense of uncertainty around the entropy values.
b. Naive Bayes classifier: A Naive Bayes classifier is a probabilistic machine learning model based on the Bayes’ theorem (Eisenstein, 2019). The Naive Bayes classifier assumes that the presence of each gesture action in the antecedent is conditionally independent given the outcome variable: it assumes that A and B are independent, given C. While this independence assumption is often violated in practice, the classifier has been shown to perform well due to its simplicity and efficiency and it is often the first classifier used for natural language processing, before exploring more complex models (Hvitfeldt & Silge, 2022). For each n-gram, the classifier calculates the posterior probability of the sequence leading to a particular next unit by multiplying the prior probability of the next unit with the conditional likelihood of the next unit given the sequence’s conditional probabilities. The outcome with the highest posterior probability is then assigned to the sequence. The Naive Bayes classifier does not encode sequence information; it learns about the probabilities at location A or B and how they affect C, but not the AB combination. Naive Bayes was implemented using the ‘tidymodels’ package ‘discrim’ in R (Hvitfeldt et al., 2023).

We also attempted to use a Long Short-Term Memory (LSTM) classifier as a type of Recurrent Neural Network architecture designed to address sequential data (Chollet et al., 2022), but faced considerable overfitting for the training data (a common issue for this type of model; Hvitfeld & Silge, 2022), so we focus on the simpler Naive Bayes classifier instead. For the Markov transitions (A->B), we used the classifier to compare the predictability of the different time-windows. For the 5-second time-window, we use the classifier to calculate how well the two antecedents independently or in combination predicted the next unit in a trigram. To reduce overfitting, we used a k-fold cross validation approach (Hvitfeldt & Silge, 2022): the dataset was cut into 50 equally-sized portions, and 98% of data were repeatedly used to predict next units in the remaining 2%. We upsampled the data to account for uneven distributions of outcomes, by randomly adding duplicates of existing cases for each outcome until all levels were equally probable (Hvitfeldt & Silge, 2022). We report prediction accuracy (correct predictions/all predictions). Within trigrams, we classified the next unit based on the correctly ordered antecedent (AB-C), the previous unit only (B-C), the first unit only (A-C), and the randomly shuffled full antecedent (BA-C). If A and B predict C equally well, we would see that units are clustered within sequences but order does not matter. If B outperforms A, we assume that there is a basic order effect based on Markov transitions (only previous unit matters). If AB outperforms B, we assume that there is some effect of combinations: adding two units together is more informative than just the previous unit. If AB outperforms B and the randomised BA, we have evidence of compositionality: knowing more than one unit and the order of both units adds information about the next unit. If AB and randomised BA perform similarly, we assume that there is an elaboration effect, where seeing both A and B together reduces uncertainty about C but that their exact order does not matter.

This comparison of a full model (AB - C) as compared with the two single-antecedent models for the same outcome data as a way to measure predictability differences of different conditions is similar to a full-null model comparison in traditional statistics and is sometimes referred to as an ‘ablation test’ in machine learning (Molnar, 2022). We use pairwise paired samples t-tests with False Discovery Rate correction (Benjamini & Hochberg, 1995) between all four conditions to determine whether some antecedents outperform others. Using model error or prediction accuracy as a measure of predictability is not without problems, as it is dependent on the model structure and complexity. Interpreting blackbox models is always difficult (Molnar, 2022), but we believe that the staggered approach chosen here allows us to gain valuable insights into a system that would otherwise be too complex for analysis, and we interpret all results in combination with the entropy measures and transition probabilities.

## Results

### Bootstrap stability

Figure 2 shows the stability of conditional probabilities at the different n-gram levels for the 5-second sequences (as the one with the largest number of transitions) by plotting the range of probabilities for each transition against the occurrence count of the antecedent unit. Figure 2.4 shows that 4-grams are essentially random, while trigrams also contain considerable uncertainty, especially for rarer antecedents. Bigrams for rare units can also take any value, meaning we have to interpret any effects with care and remove any rare combinations.

### First Order Transitions Across Time-Windows

#### Significant N-grams

For bigrams, the four time-windows differed considerably in the number of transitions that occurred significantly more than expected at an alpha 0.05 level and that occurred at least 5 times. Longer time-windows produced more significant transitions than shorter time-windows. For the 5-second sequences, 101 out of 476 transitions that were observed in total (21.2%) were significantly more likely than expected under the null model. For the rapid-fire sequences, 57 out of 315 observed transitions (18.1%) were significant. For the overlap sequences, 35 out of 187 observed transitions (18.7%) were significant; for the solitary-gestures-with-waiting sequences, 30 out of 242 transitions (12.3%). There were no transitions that were significant in any of the shorter time-windows that were not also significant in the longest time-window (5-seconds). We found very few indications of deterministic transitions. The only deterministic transition (conditional probability equals 1) occurred in the overlap network, where all Embrace gestures were followed by Bite gestures (but the same bigram was not deterministic in the longer time-windows). The time-windows differed in the number of observed repetitions; in the solitary-gesture-with-waiting sequences, 45% of gestures in sequences were followed by the same gesture, compared to 18% in overlap sequences (because this would necessitate doing the same gesture with different limbs), with the other time-windows (rapid-fire sequences: 27%; 5-second sequences: 34%) in between.

#### Transition Network Structure

We present the transition network based on the concurrence of the different time-windows - edges are included if they are significant in at least 3 out of 4 time-windows, to ensure that the presented network represents one that is largely independent from researcher choices of sequence definition (Fig. 3). In the Supplementary, we present a table of all significant transitions for all time-windows. The transition network revealed five clusters containing more than one repeated unit. These were clustered around modality (contact vs non-contact vs object-contact) and context; e.g., one larger play initiation cluster, one grooming initiation cluster, and several for redirecting partner attention. One cluster consisting of *GrabHold*, *Grab*, *Bite*, and *Embrace* were contact gestures that allow individuals to hold the partner in close proximity. *Big Loud Scratch*, *Present*, and *Raise* are all closely associated with grooming requests. This grooming cluster is connected to a cluster associated with repositioning or reorienting the partner (*Push* and *Touch*, with the latter also being a contact extension of *Reach* in a begging context). *Dangle*, *Swing* and *Object Shake* are all gestures performed while holding a static object. The largest cluster (*HitObject, StompObject, ObjectMove, ObjectShake, Jump*) contains object manipulations that can act as play invitations.

**Figure 3:**
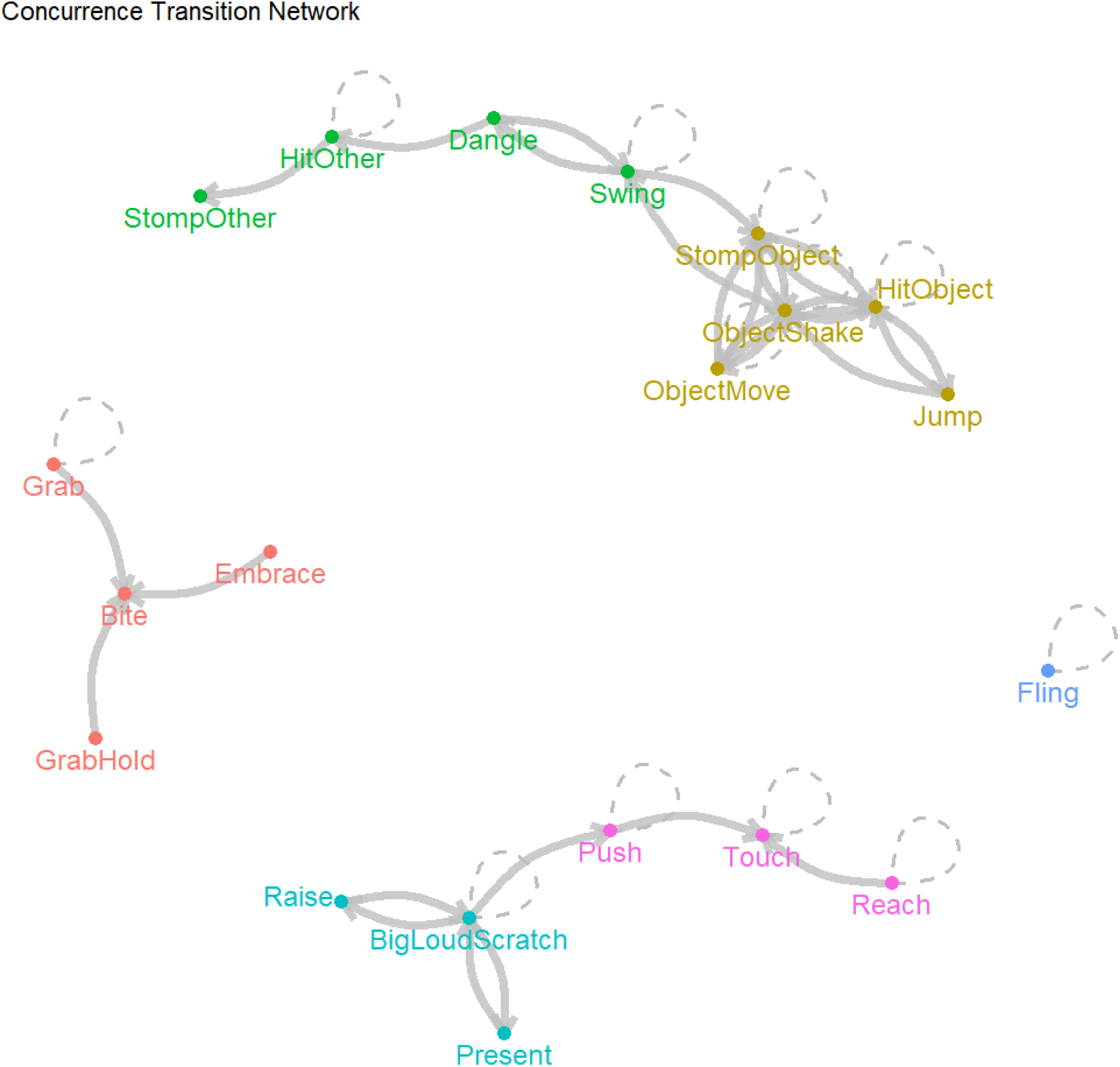
Transition network based on the concurrence of the different time-windows (i.e., transitions that are significant in at least three out of four time-windows) including cluster assignment based on ‘optimal clustering’ algorithm and loops.

When comparing the similarity of which transitions were significant in the different time-windows, the two sequence definitions with response waiting of below 1 second between individual units (rapid-fire sequences, overlap) produced highly similar results. Figure 4 shows transitions that were the same in the solitary-gestures-with-waiting and rapid-fire sequence networks (left), those that were significant only for the rapid-fire sequences (middle), and those that were significant only after solitary gestures with response waiting (right).

**Figure 4:**
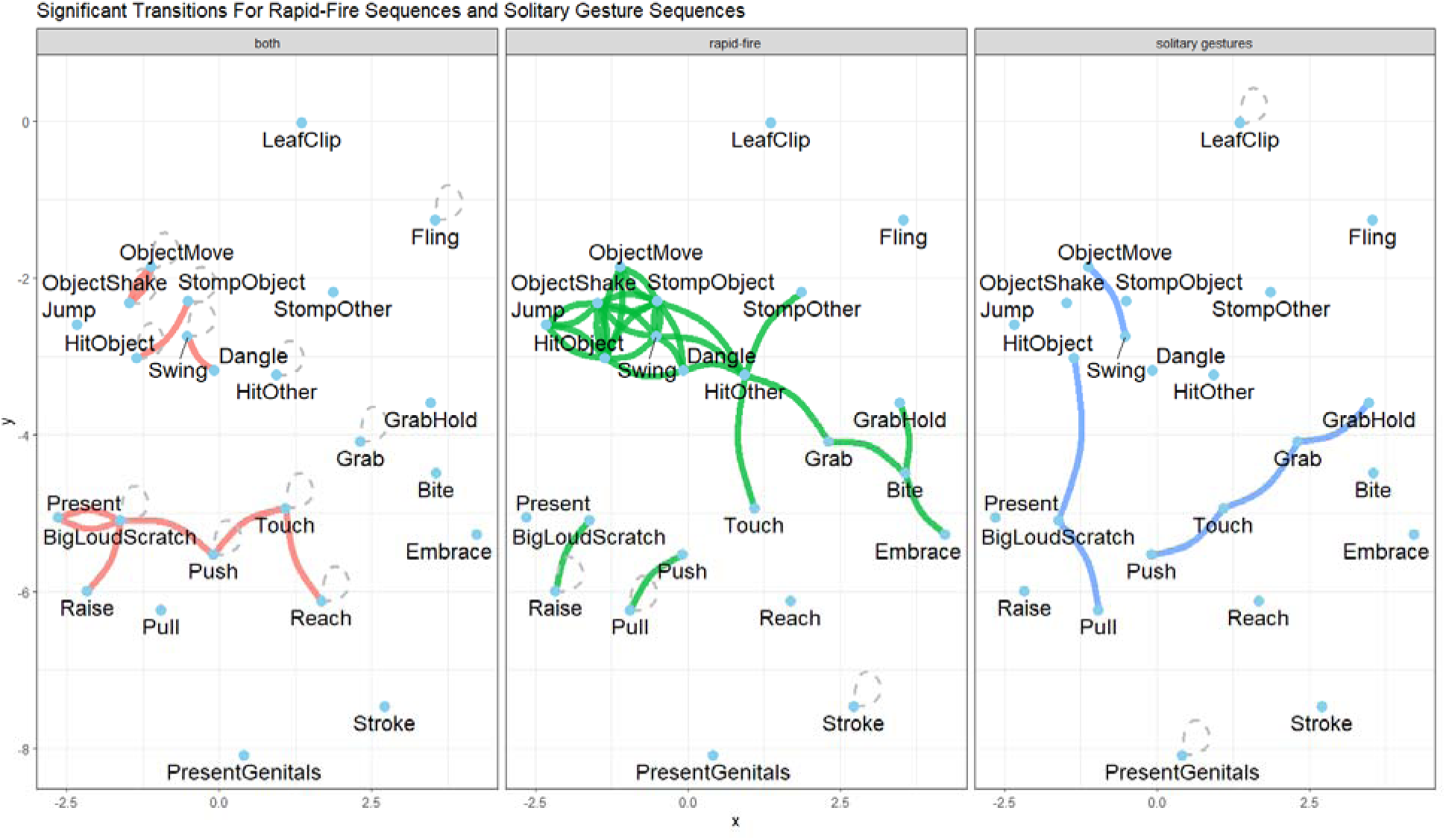
Network of significant transitions that are shared between the rapid-fire sequences and solitary gesture sequences (left, red), occur only in the rapid-fire sequences (centre, green) or only in the solitary-gestures-with-waiting sequences (right, blue). Significant transitions are marked by edges between gesture actions. Loops indicate that a gesture action significantly follows itself.

From Figure 4, we see that those transitions that were specific to rapid-fire sequences were primarily focused on object-related gestures and those that can be achieved using different body parts (e.g., *Bite* and *Embrace/Grab*). Solitary-gestures-with-waiting sequences more commonly included gestures that would not be marked as distinct if the response waiting was not observed (e.g., *Grab* and *GrabHold*, or *Touch* and *Push* would probably be marked as one of the options without the behavioural indication of waiting separating them).

#### Bigram order

There were 21 cases where AB was significantly more common than predicted for BA given random distribution within the 5-seconds sequences, see Table 3. Those were usually related to contact gesture actions, which tended to follow non-contact gesture actions. Aside from the 7 most directional combinations, in which the conditional probability was more than 5% higher than expected, effect sizes (measured as the difference between the observed and expected conditional probability) here were small. The other time-windows follow the same pattern (with reduced numbers of cases because of smaller sample sizes) with the exception of solitary gesture sequences, in which the effect sizes (how much more likely than expected was a transition) were even smaller, indicating that instances of gesturing clearly differentiated by a period of response waiting did not show order effects.

**Table 3:** Bigrams where one unit was more commonly observed to precede the other than would be expected given random distribution within sequences, arranged by the strength of effect. For the 5-second time frame only.

| Antecedent | Consequent | Count | Conditional probability | Expected | Improvement |
| --- | --- | --- | --- | --- | --- |
| Embrace | Bite | 16 | 0.64 | 0.39 | 0.25 |
| GrabHold | Bite | 18 | 0.42 | 0.23 | 0.19 |
| Grab | Bite | 18 | 0.25 | 0.13 | 0.12 |
| Bite | HitOther | 6 | 0.17 | 0.1 | 0.08 |
| Raise | BigLoudScratch | 23 | 0.34 | 0.28 | 0.07 |
| Push | Touch | 15 | 0.16 | 0.11 | 0.05 |
| Raise | StompObject | 5 | 0.07 | 0.04 | 0.04 |
| Pull | Push | 7 | 0.11 | 0.07 | 0.04 |
| Swing | HitOther | 11 | 0.09 | 0.06 | 0.03 |
| ObjectShake | StompObject | 35 | 0.13 | 0.1 | 0.03 |
| HitOther | StompOther | 8 | 0.06 | 0.03 | 0.03 |
| Push | Raise | 6 | 0.07 | 0.04 | 0.03 |
| BigLoudScratch | Push | 14 | 0.05 | 0.03 | 0.02 |
| HitObject | Jump | 9 | 0.04 | 0.02 | 0.02 |
| ObjectMove | ThrowObject | 5 | 0.04 | 0.02 | 0.02 |
| Touch | HitOther | 5 | 0.04 | 0.02 | 0.02 |
| Touch | GrabHold | 7 | 0.05 | 0.04 | 0.02 |
| StompObject | HeadStand | 5 | 0.02 | 0.01 | 0.02 |
| BigLoudScratch | PresentGenitals | 5 | 0.02 | 0.01 | 0.01 |

#### Entropy

In this analysis, we compared the information content (measured in bits) of conditional probabilities at different levels with what would be expected given random distribution of units, represented by the ratio of the two. Because of the sample size differences for the four time-windows, we present a graph that shows the entropy of the first-order transition for different subsamples of each time-window (Figure 5). The change in entropy with increasing sample size was similar for the different time-windows, with all of them showing similar ratios of observed and expected entropy (around 0.5-0.6) already with relatively small samples. An entropy ratio of 0.5 indicates that the observed entropy is half the expected entropy under the null model of random distribution of units, a sign of a mix of predictability and unpredictability at this level. The overlap sequences and sequences containing solitary-gestures-with-waiting, as the most restrictive definitions of what constituted a sequence, showed lower entropy than the other time-windows, but the effect was small. Table 4 contains the observed and expected entropies for the full data for all time-windows.

**Figure 5:**
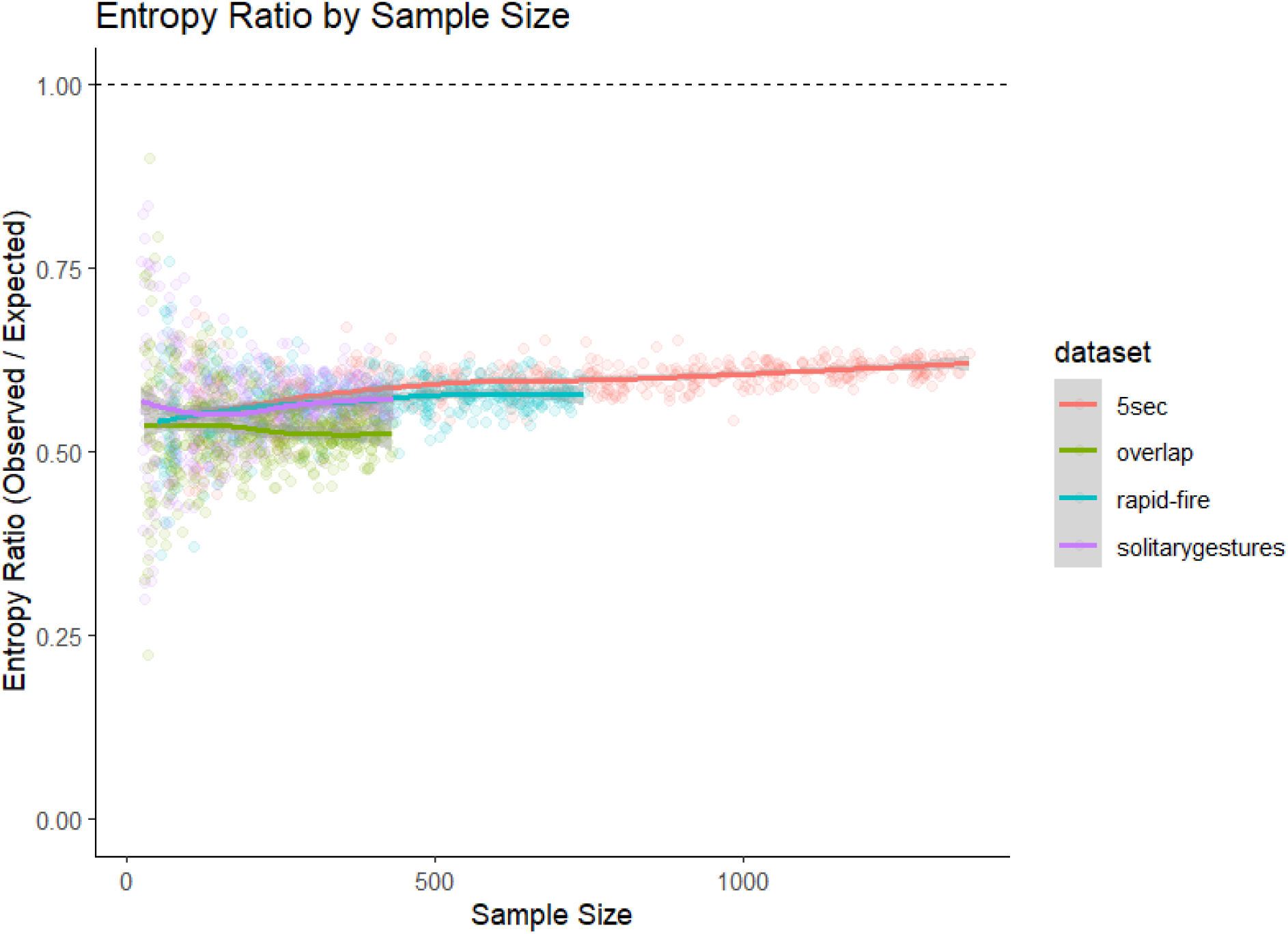
Entropy ratio values of conditional probabilities with antecedents of one unit. Colours indicate the time-window. Entropy ratios (observed/expected entropy) closer to 0 are considered more predictable than random assignment, 1 or above are random distributions.

**Table 4:**
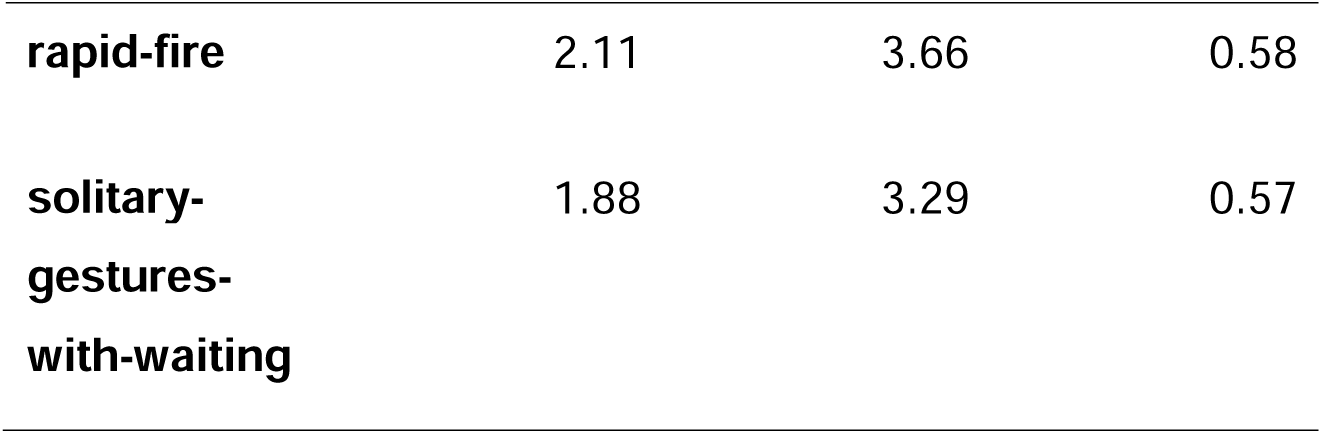
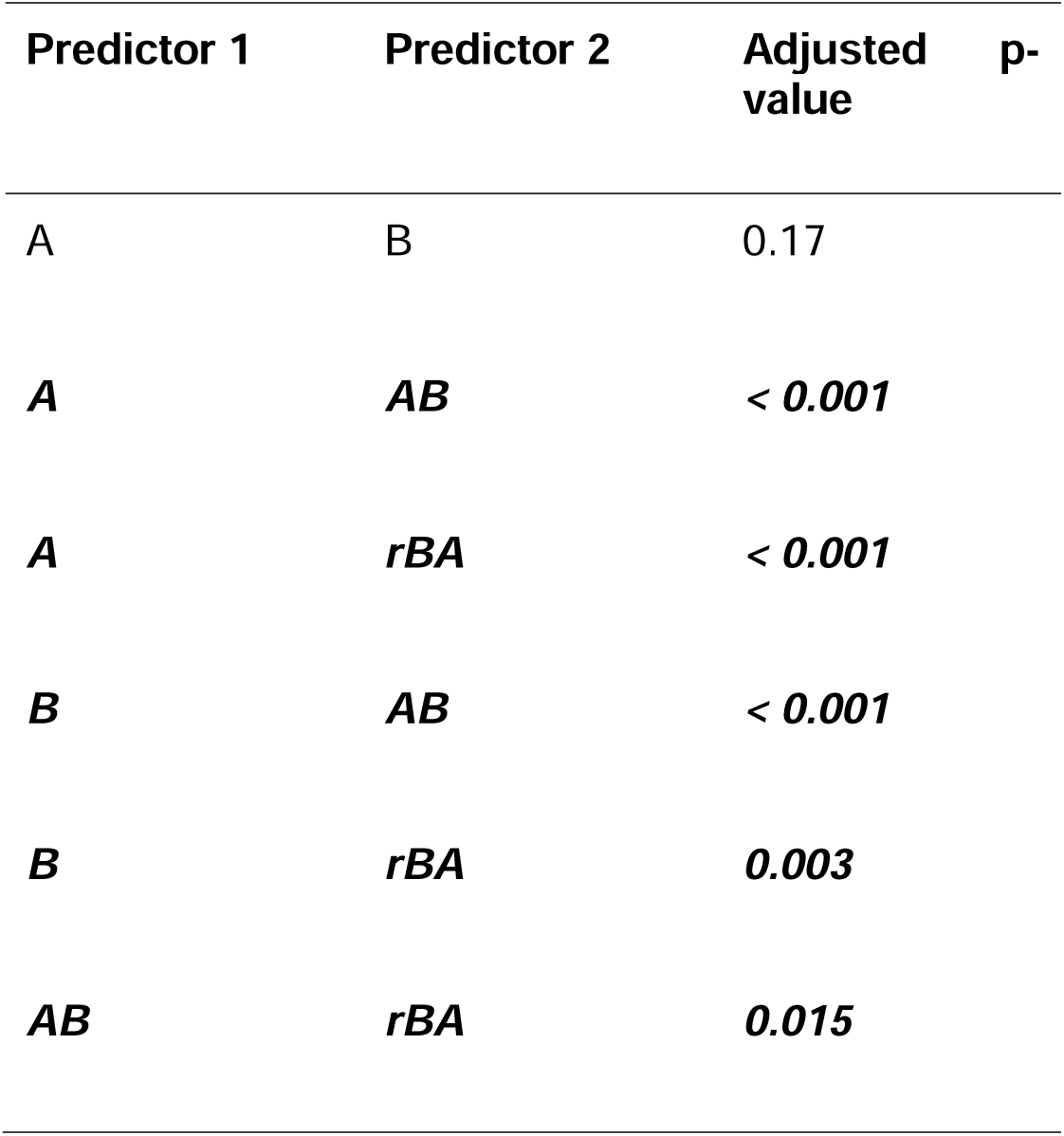
Observed and expected entropies for the four time-windows, considering the full dataset for each.

**Table 4:**
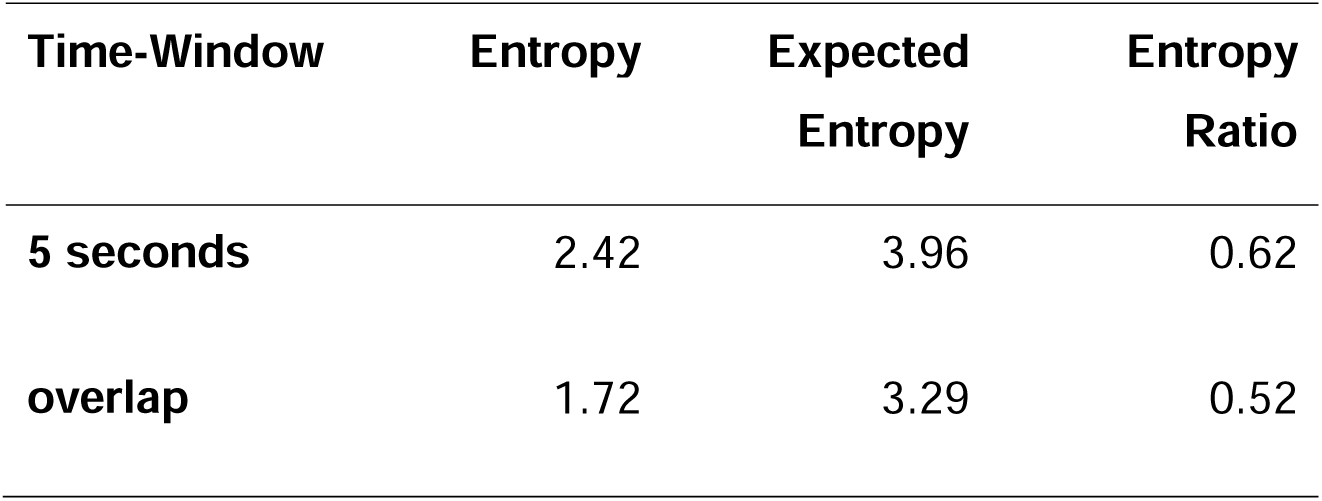
Results of pairwise comparisons of prediction performances (measured using prediction accuracy) of Naive Bayes classifier using paired samples t-tests with False Discovery Rate correction of p-values. Significantly different pairings marked in italics bold.

#### Next element prediction

In this analysis, we first compared the prediction accuracy for the four time-windows, assuming first-order transitions. Figure 6 shows the prediction accuracy of k-fold cross validation with k = 50 using the Naive Bayes classifier (i.e., 98% of the data is used for training data), applied to subsets of each time-window of different sample sizes to account for the differences between datasets. We observe that increased sample sizes improved predictions, but that this trend was not uniform. It was very pronounced for the solitary gesture sequences, which also had the highest prediction accuracy despite the smallest overall sample size (accuracy = 0.31), probably because of a high level of repetitions. The other time-windows had accuracies that were comparable to each other for similar sample sizes (5-seconds: 0.29; overlap: 0.24; rapid-fire: 0.21). That means that even without further information about the context or individuals or other modalities, a basic classifier with a small training set could achieve almost 30% correct classifications of the next unit in a sequence based on basic occurrence and transition probabilities.

**Figure 6:**
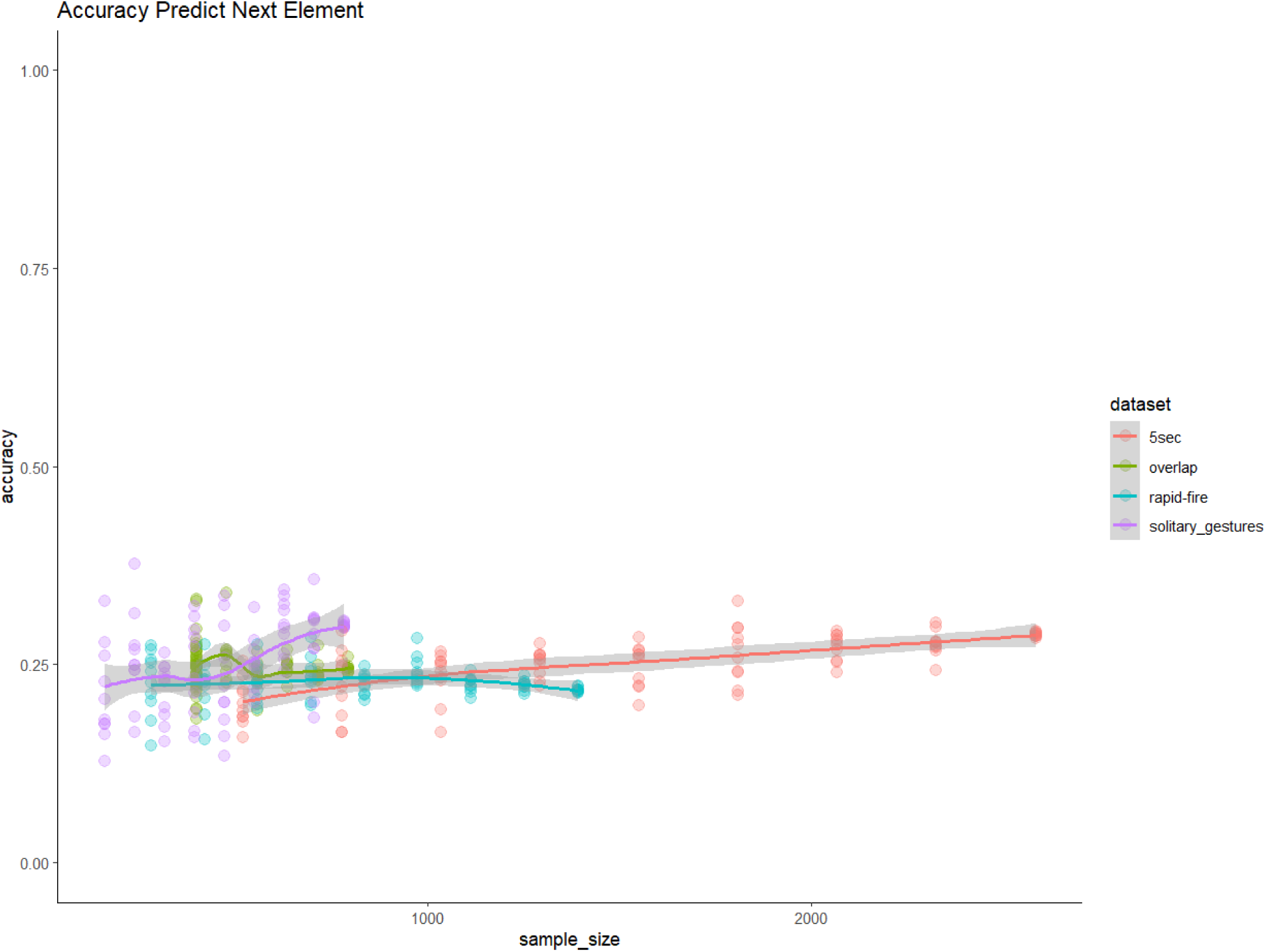
Accuracy values of Naive Bayes classifier (correct predictions/all predictions), predicting the consequent units based on antecedents of one unit. Colours indicate the time-window.

### Second Order Transitions For 5-Seconds Sequences

#### Significant N-Grams

For the 5-seconds sequences and focusing on trigrams, 31 out of 593 transitions that were ever observed were significant and occurred over 5 times (see Supplementary Material). All significant trigrams contained at least one occurrence of the consequent in the antecedent (ABA or BAA), and often both antecedent units were the same as the consequent (AAA). All significant antecedents in the trigrams were also significant bigrams.

#### Entropy

For the 5-seconds sequences, we could also analyse trigrams, including the entropy under the assumption that order does not matter (by alphabetising antecedents before calculating probabilities, so that AB and BA were considered together). Figure 7 shows that at the bigram level (one antecedent known), the observed entropy was lower than would be expected, indicating that the observed transition probabilities indeed reduced uncertainty about which unit was seen next. For trigrams, this effect was less pronounced, with a higher entropy ratio when the order of antecedent units was considered; probably due to the small sample size for most trigrams. However, when removing the order information from the antecedent (thereby also increasing the sample for each distinct antecedent), we saw that knowing the antecedent reduces the entropy as compared to the random distribution. This outcome indicates that, at least at the sample sizes observed here, order adds noise rather than reducing it.

**Figure 7:**
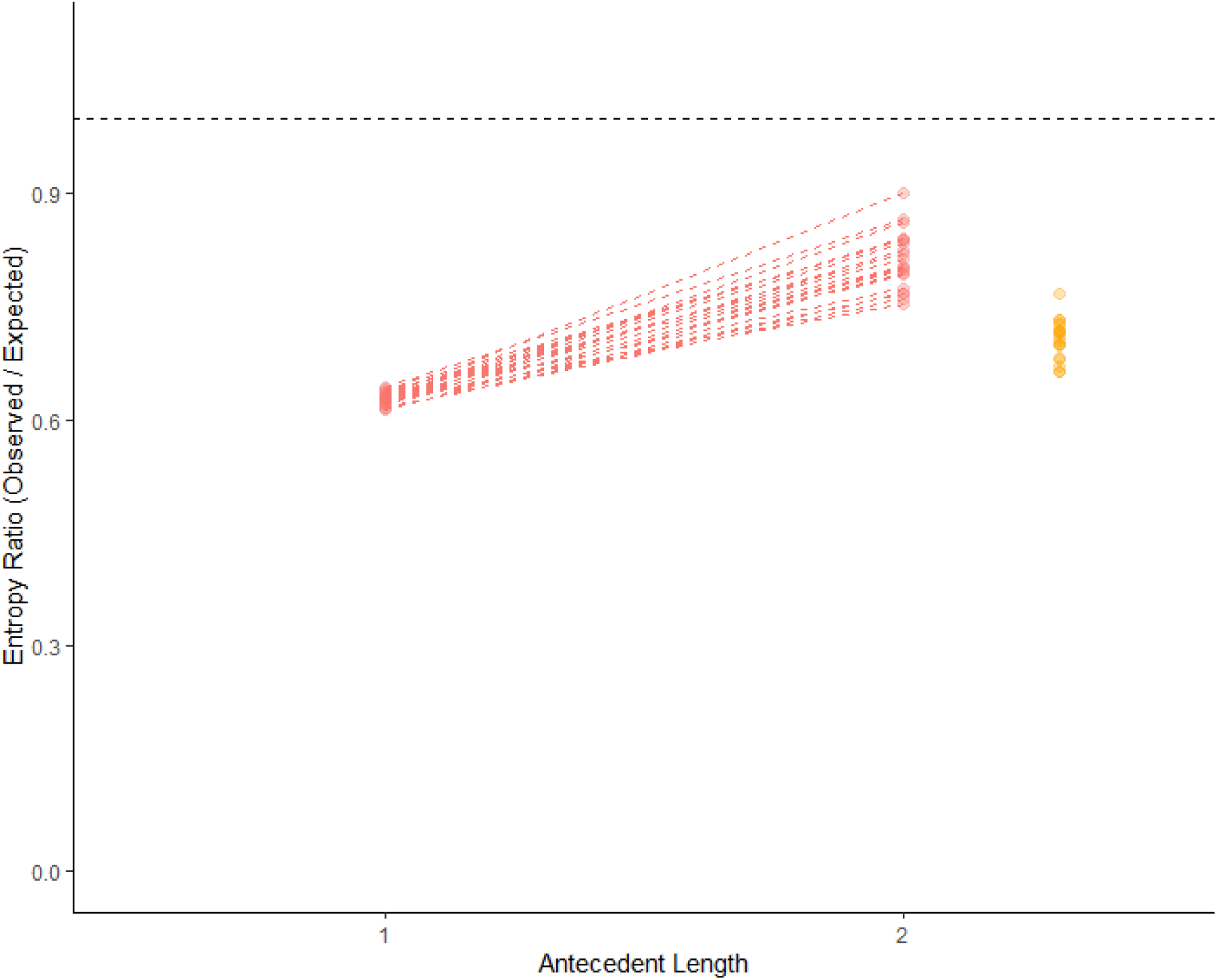
Entropy values of conditional probabilities with antecedents of 1 or 2 units. True order in red. Each line represents one bootstrap (resampling with replacement) to indicate uncertainty of estimates. Values closer to 0 mean that observed values are lower than expected values and the system is more ordered and more predictable than random assignment. ‘Alphabetical’ ordering for 2-unit antecedents (portrayed in orange) means that the constituting units were ordered alphabetically before calculating probabilities, removing order information.

#### Next Element Prediction

We compare the prediction accuracies of models containing only the unit two steps removed from the consequent (A), the immediately preceding unit (B), both units together (AB), and both units together with order information removed through alphabetising them (rAB). As a reminder, if A outperforms B, then we assume Markov effects. If AB outperforms B alone, we count this as evidence for possible composition effects. If AB and rAB perform similarly, we consider this evidence for an elaboration effect, where order does not matter but having two gestures reduces uncertainty about the subsequent order. If AB outperforms rAB, we consider that order matters for the purposes of prediction.

In Figure 8 and Table 4, we see that B (accuracy: 0.185) does not significantly outperform A (accuracy: 0.17). AB (0.27) and rAB (0.25) performed better than A or B alone, and AB slightly outperforms rAB.

**Figure 8:**
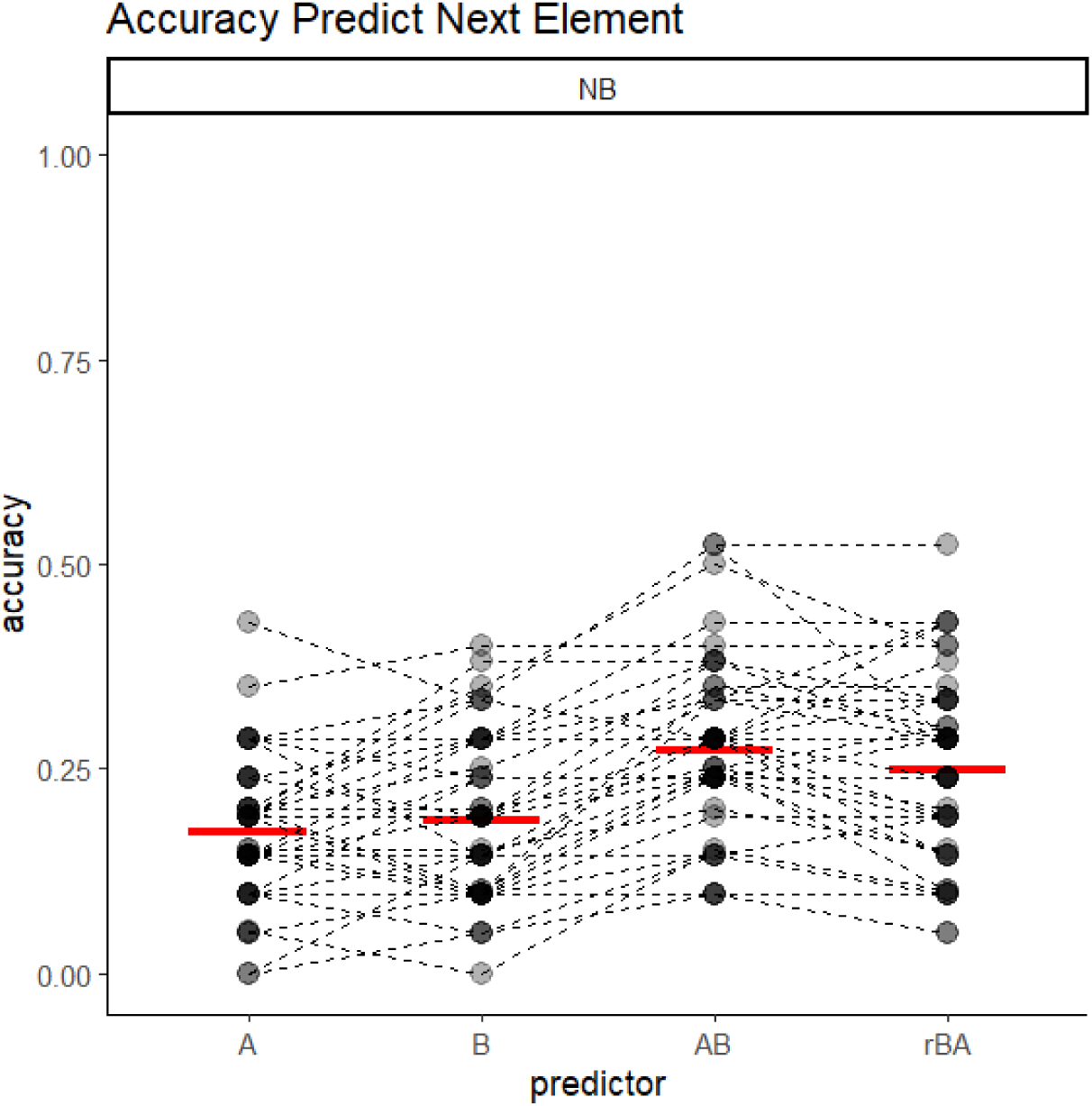
Correct classification probability of models using Naive Bayes classifier. Lines connect iterations based on the same training/test dataset split. Red lines indicate average performances of classifiers. Prediction based on two units removed (A), previous unit (B), two preceding units (AB), and two preceding units without order information (rAB).

When testing which gesture actions were more accurately classified based on the two preceding units, rather than on one preceding unit alone, Figure 9 shows that there are a small number of units where the single predictor performed more accurately (e.g., *Head Stand*, *Throw Object*, *Stomp Other*, and *Present*); some where neither antecedent had any predictive power; some showed the same performance (these were all very rare); some where the single unit had no predictive power but the combination performed quite well; and some where both showed some accuracy but accounting for more antecedent units improved performance.

**Figure 9:**
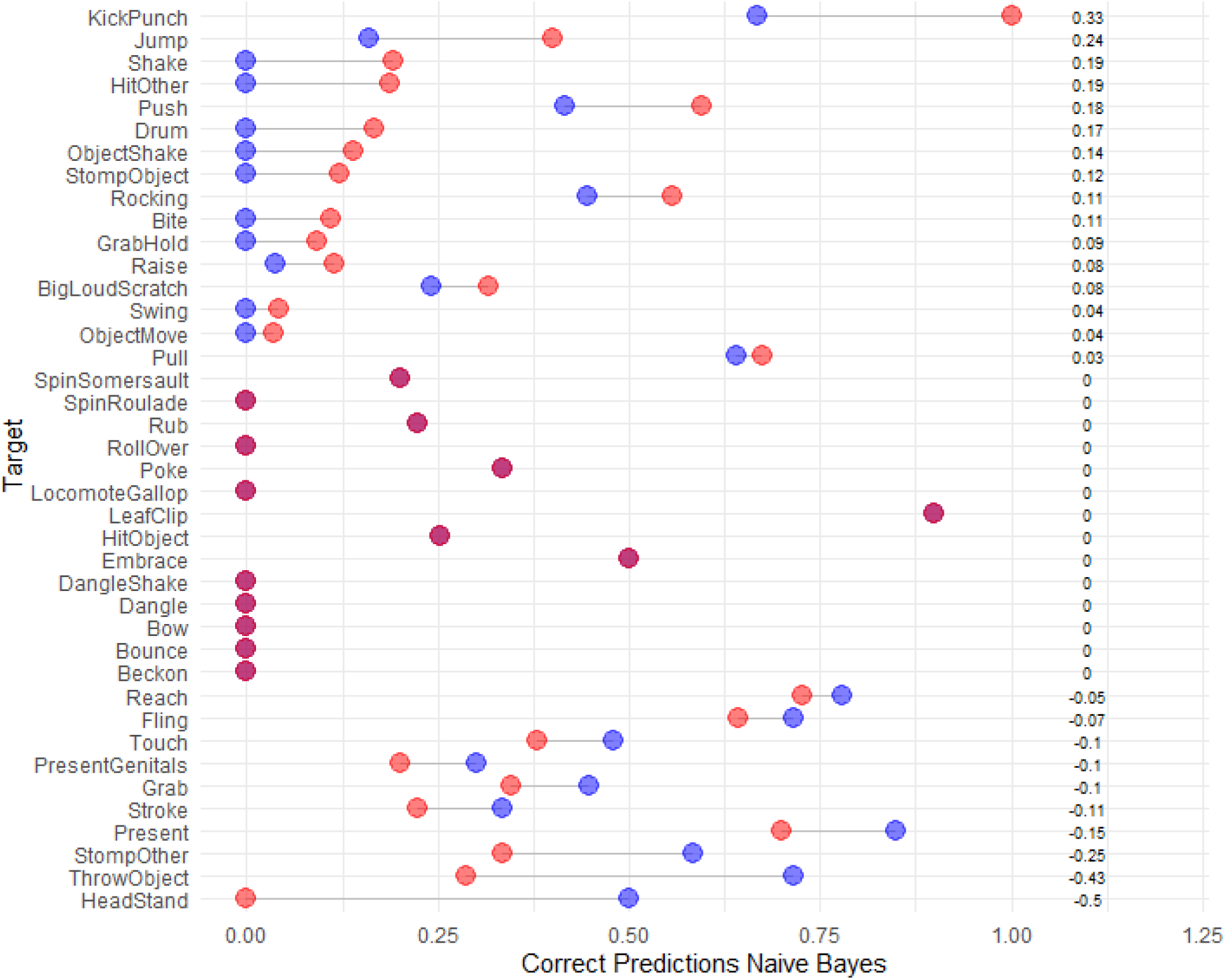
Comparison of unit-level prediction accuracy for the Naive Bayes classifier for each gesture - prediction based on the single antecedent B in blue, for the two-antecedent AB combination in red, with the difference on the right. Where only one dot is visible, the predictive power was the same.

## Discussion

In this study, we analysed transition patterns in Eastern chimpanzee gesture sequences. Combinatorial capacities in naturally occurring communication sequences have recently been reported in a variety of species, but with a strong focus on vocalisations and song-like communication (Allen et al., 2019). For chimpanzees, reliable associations between units have been described in vocal (Girard-Buttoz et al., 2022; Leroux et al., 2021) and multimodal (Oña et al., 2019) sequences. Here, we showed that sequences in the chimpanzee gestural system, characterised by a larger number of meaning-bearing and intentional signals, were generally short and there were many rare units and rare transitions between units that did not pass our thresholds, mirroring earlier analyses with a subset of the current dataset (Hobaiter & Byrne, 2011a).

We first asked whether there were non-random transitions between elements. We identified a number of transitions that occurred at higher-than-expected frequencies, with sample sizes limiting us to statements about combinations of one or two antecedent units (and considerable variation for the latter). Around one third of the 101 significant first-order transitions in a 5-second time-window were repetitions of the same unit, indicating that persistence or redundancy constitute an important part of chimpanzee gesture sequences. This outcome is expected, given that intentionality is based on persistence as a core property (Bates et al., 1975; Leavens et al., 2005), so what is considered a ‘gesture’ here (as compared to other actions) is biased towards repetitions. In both earlier studies (Hobaiter & Byrne, 2011b) and in our own findings the likelihood of a gestural unit being repeated increased strongly after a period of response waiting.

When combining all time-windows, we identified a core network of transitions between units that could be detected independently of the time-window in question. For some element pairs, only one direction was significantly more likely than expected, potentially because of physical constraints of actions: for example, individuals *Grab*, *GrabHold*, or *Embrace* a partner (establishing sustained physical contact) and then *Bite* them; they would also first *Reach* and then *Touch*, and first *Dangle* from a branch, then *HitOther*, then *StompOther.* In other cases, there was a higher probability that one action precedes, but both directions had significantly higher probabilities than chance, for example: *Present*, *BigLoudScratch*, and *Raise* (all common in grooming initiations) were observed in any bigram order, even though *BigLoudScratch* usually followed the *Raise.* Observed clusters of significant transitions were a mix of repetitions (loops in the network), physical contingencies (e.g., object-related gestures co-occurring at high rates), potentially synonymous actions (e.g., *Grab*, *GrabHold*, and *Embrace* preceding the same gesture), and shared meaning (*Present*, *BigLoudScratch*, and *Raise* as grooming gestures).

Did unit order matter? We identified a small number of two-element dyads where transition direction seemed to significantly matter, but they either had very small effect sizes (probability increase for one direction over that expected of 5% or below) or the order effect could be explained with relative ease through physical affordances (individuals first grab a partner to pull them closer, then bite them). In the entropy measures, the correct order was noise that increased entropy as compared to the alphabetised order (indicating that it was the co-occurrence that mattered, not the production order). For the prediction models, the correctly ordered combination performed better than the alphabetised order, low level evidence that order information improved predictability. Thus, it appears as if combinations of gestures serve to disambiguate the subsequent elements in a system in which there are very large numbers of units, each associated with different sets of goals (Graham et al., 2018, 2020). For example, *Big Loud Scratch*, a very common gesture for which we can be fairly certain about the transition probabilities, was followed by 15 different gestures when considered by itself, but only by three when it was associated with *Present*; mainly by repetitions of *Present* or *Big Loud Scratch*. However, *Big Loud Scratch*/*Present* and *Present*/*Big Loud Scratch* did not lead to different consequent units, showing either that order did not matter for next-unit prediction or that applying strict linear temporal structure to the definition of gesture sequences was insufficient. Unlike in most vocalisations, multiple gesture actions can be performed simultaneously and gestures may not be restricted to the linear ordered structure that we assume for animal vocal communication. Other non-vocal communication systems, such as signed languages (Armstrong et al., 1994; Sandler & Lillo-Martin, 2006; Stokoe, 1980), incorporate spatial dimensions into their syntactic structure, but it remains speculative whether the same is true for chimpanzee gestures.

Regarding the predictability of the system as a whole, information about previous gestures reduced uncertainty about consequent gestures, with two antecedent gestures improving prediction accuracy and entropy ratios as compared to a single one, similarly to the pattern found in a recent study of marmoset call combinations (Bosshard et al., 2024). We found conflicting evidence on whether the order of the antecedent units mattered: alphabetising the order of AB in a bigram antecedent led to worse predictions, but reduced the entropy ratio. This result could be explained by methodological differences; the Naive Bayes classifier uses the joint conditional probabilities of the two antecedents in their position predicting the consequent, while the entropy uses the conditional probability of the combination itself while disregarding the probability of each individual antecedent. At the same time, we found a remarkable level of predictability: even a basic classifier like Naive Bayes, using the relatively small datasets available (as compared to typical machine learning datasets), achieved prediction accuracy of around 30% for bigrams, well above chance. Encouragingly, bigram entropies could be established accurately with relatively small sample sizes for all time windows and were remarkably consistent across the different datasets. This level of robustness, indicates that the choice of time window had only minor impacts on overall predictability of the resulting sequence system, and that system-level parameters are less dependent on researcher choices than transition patterns. It also suggests that this approach would be reliable in the smaller datasets typically available for most studies of ape gesture. In terms of predictability, the two sequence definitions with the most restrictive rules (i.e., gestures have to overlap or gestures have to be separated by 1 second) were the most predictable using both entropy and classification accuracy, because of their much smaller transition networks.

Comparing different time-windows for the bigram transitions revealed an impact of methodological decisions, with potential biological implications. Given different traditions within the field for defining what constitutes a sequence based on temporal and/or behavioural criteria (with a mix of different a priori or data-driven time thresholds and the inclusion or exclusion of sender and receiver behaviour in delineating sequences), this is potentially worrisome. In many cases the investigation of group- or species-level comparisons in ape signalling is reliant on comparison across samples and studies where methodological decisions are not necessarily similar or transparent (Rodrigues et al., 2021). Our findings suggest that direct comparisons between studies that involve different approaches could be problematically biased. As in any signal detection task, we face a sensitivity-specificity trade-off: reducing restrictions (by considering larger time-windows) increases the sample size and reduces false negative results, but potentially leads to more false positives and the inclusion of noise, for example: different gesture sequences with different goals being grouped together for longer thresholds. Using a time-based approach based purely on gesture start and end times (using 5 seconds from the end of the previous MAU) created more and longer sequences than an approach based on response waiting or overlap, with considerably more significant transitions between units. Given sample size and time constraints in many studies, creating sequences based on temporal thresholds without regard for sender or recipient behaviour will remain an important approach for many researchers. Reassuringly, all transitions that were identified as significant in either of the more restrictive time-windows were also represented in this largest time-window. Using a time-based approach with 1 second intervals was nearly indistinguishable from the rapid-fire sequences and was therefore dropped, indicating that researchers who do not code behavioural markers to distinguish sequences can nevertheless rely on time-based approaches and get similar results. Approaches that use strong a priori reasoning for restricting the time-window (e.g., based on recipient behavioural markers) benefit from higher resolution and potentially allow researchers to differentiate structural dimensions of sequences. In our case, the difference between the two most distinct time-windows (rapid-fire and solitary gestures) could partially be explained by the inability for coders to distinguish certain gesture types if they were to occur in rapid sequence. In the absence of strong data- or theory-driven thresholds or markers, we advocate the use of multiverse analysis. Doing so allows studies using different thresholds that are not directly comparable to include multiple thresholds and report outcomes, increasing replicability and reuse (Hoffmann et al., 2021; Mielke, 2023; Steegen et al., 2016).

In comparison to vocal sequences in Western chimpanzees (Girard-Buttoz et al., 2022), gesture sequences were made up of considerably more independent units with much lower occurrence frequencies, partially as a result of methodological decisions to lump vocalisations, making direct comparisons difficult. For example, the least common unit in the vocal study occurred 78 times, which is more common than half of our gesture actions, impacting the reliability of the calculated conditional probabilities. However, following the definitions provided by Girard-Buttoz et al. (2022), the gesture system showed high levels of flexibility (each unit is followed by multiple others); limited ordering effects (each unit is associated with a small subset of other units, with a few cases of non-random order within bigrams); and recombination (two antecedent units improve prediction accuracy for subsequent units over single units, indicating that they are repeatedly used with additional units in sequences). In the latter case, the strongest rule for trigrams was repetition: all significant two-unit antecedent combinations contained the consequent as either one or both antecedent units; sequences often contain redundant information or persistence, perhaps in response to recipients who do not react in the expected or desired way. Overall, we remain cautious when comparing sequences of different signal types as these may stem from repertoires determined using different criteria and thresholds, which in turn may affect sequence number and diversity. We encourage researchers working on different species and signals to apply transparent methods and find common ground that allows more comparable approaches.

More and more animal species and communication systems show rudimentary combinatorial capacities, with some indications of basic syntactic structures (Berwick et al., 2011). Largely confirming previous results on great ape gestures (Genty & Byrne, 2010; Graham et al., 2020; Hobaiter & Byrne, 2011a; Liebal et al., 2004) and mirroring vocal research (Leroux et al., 2021) using a larger dataset, we showed that chimpanzee use short sequences of gestures, and that transitions in the sequences are not randomly distributed. Significant transition patterns seemed to be driven by a mix of affordances (e.g., performing different actions with an object) and shared meaning (e.g., grooming or play initiations). We found that these rules make the system more predictable. At the same time, while we found biologically-relevant order-effects, these were most likely shaped by the ease of signal production, indicating that for the majority of sequences linear-ordering was unlikely to shape information. This result does not preclude compositional or syntax-like structure, but indicates that a more targeted approach than repertoire-wide transition probabilities is necessary to move from combination to composition. The combination of elements in sequence to expand the information-bearing capacity of a system is far more likely to be relevant in smaller sets of signal units. The chimpanzee gesture system is currently estimated to contain somewhere between 70 and 140 units (Mielke et al., 2024), potentially an order of magnitude greater than their vocal repertoire. The information content of meaning-bearing ape gestures is flexible — with most units employed to express 2-3 meanings (Graham et al., 2018; Hobaiter & Byrne, 2014). As a result, the structured combination of signals in gestural sequences may serve to disambiguate meaning in a specific *occasion* of use. The ability for the same signal to encode different meanings on different occasions has been suggested as a critical component of language use. As experimental studies of gestural communication are near impossible, incorporating meaning into observational analyses of sequence production will provide a valuable next step to the approach taken here. Formalising the syntactic structures present in signed languages required linguists to reconsider the nature of the structural dimensions they described (Stokoe, 1980), fully describing the combinatorial properties of ape gesture may require a similar reimagining of the ways in which non-human gestural systems encode information.

## Acknowledgements

AM was funded by a Leverhulme Early Career Fellowship. CH, GB, KEG, and AS were supported by funding from the European Research Council under Gestural Origins Grant No: 802719. KS and CW were supported by funding from the European Research Council under Grant No: ERC_CoG 2016_724608. We thank all the staff of the Budongo Conservation Field Station, its founder Vernon Reynolds, and the Royal Zoological Society of Scotland who provide core funding. We thank the directors of the Kibale Chimpanzee Project for permission to use video data archives. We thank the Uganda Wildlife Authority, the National Forestry Authority, the President’s Office, and the Uganda National Council for Science and Technology for providing research permits and permissions to conduct research in Budongo, Kalinzu, and Kanyawara. The Issa project (GMERC) is grateful for long-term support provided from the UCSD/Salk Center for Academic Research and Training in Anthropogeny (CARTA). We thank the Tanzanian Wildlife Research Institute (TAWIRI), Commission for Science and Technology (COSTECH), and Tanganyika District for permission to conduct research in the Issa Valley. We thank all the field assistants and local staff across field sites for the decades of work that make this kind of research possible.

## Data availability

All scripts and data can be found here: https://github.com/AlexMielke1988/Mielke-et-al-Ngrams. Data are provided as prepared sequences only to allow for replication; for the full data, please contact the first or last author.

