## Supplementary for "Predictability of next elements in chimpanzee gesture sequences"

***Table S1: Gesture actions – Pre-defined lumping decisions and frequencies***

| **Gesture action** | **Lumped with** | **Frequency** | **Modifiers with variation** |
| --- | --- | --- | --- |
| Beckon | - | 32 | Body_part_signaller |
| Big Loud Scratch | - | 871 | Body_part_contact, Body_part_signaller, Laterality |
| Bite | Bite kiss, Bite threat | 163 | Body_part_contact |
| Bounce | - | 14 | - |
| Bow | - | 12 | - |
| Dangle | - | 241 | - |
| Dangle Shake | - | 40 | Repetition |
| Drum | - | 23 | Body_part_signaller, Laterality, Repetition |
| Embrace | - | 88 | Body_part_contact, Body_part_signaller, Laterality |
| Fling | - | 100 | Body_part_signaller |
| Grab | - | 205 | Body_part_contact, Body_part_signaller, Laterality |
| Grab hold | - | 112 | Body_part_contact, Body_part_signaller, Laterality |
| Head stand | - | 38 | Body_part_contact |
| Hit object | Hit Object Object, Hit Object Soft, Hitting Object, Hitting Object Object, Hitting Object Soft | 484 | Body_part_contact, Body_part_signaller, Laterality, Repetition |
| Hit other | Hit Other, Hit Object Other, Hit Other Soft, Hitting Other, Hitting Other Clap, Hitting Other Soft | 322 | Body_part_contact, Body_part_signaller, Laterality, Repetition |
| Jump | - | 38 | - |
| Kick punch | Kick punch at | 10 | Body_part_contact, Body_part_signaller |
| Leaf clip | Leaf Clip Drop | 54 | Body_part_signaller, Laterality, Repetition |
| Locomote gallop | - | 49 | - |
| Lunge | - | 13 | - |
| Object mouth | Object Mouth Attached, Object Mouth Unattached | 31 | - |
| Object move | Object Move Attached, Object Move Unattached, Rake object | 256 | Body_part_signaller, Laterality, Repetition, |
| Object shake | - | 589 | Body_part_signaller, Laterality, Repetition, |
| Poke | Poking | 12 | Body_part_contact, Repetition |
| Present | Present directed | 661 | Body_part_signaller, Laterality |
| Present genitals | Present genitals forwards, Present genitals backwards | 345 | Body_part_contact, Body_part_signaller |
| Pull | Pull directed | 188 | Body_part_contact, Body_part_signaller, Laterality |
| Push | Push directed | 399 | Body_part_contact, Body_part_signaller, Laterality |
| Raise | - | 236 | Body_part_signaller, Laterality |
| Reach | - | 587 | Body_part_signaller, Laterality |
| Rocking | Rocking Bipedal, Rocking Sit | 36 | - |
| Roll over | - | 34 | - |
| Rub | - | 59 | Body_part_contact, Body_part_signaller, Repetition |
| Shake | - | 77 | Body_part_signaller, Laterality, Repetition |
| Spin roulade | - | 23 | - |
| Spin somersault | - | 42 | - |
| Stomp object | Stomping object | 425 | Body_part_signaller, Laterality, Repetition, |
| Stomp other | Stomping other | 31 | Body_part_contact, Body_part_signaller, Laterality, Repetition, |
| Stroke | - | 40 | Body_part_contact, Body_part_signaller, Laterality, Repetition |
| Swing | Swing directed | 279 | Body_part_signaller, Laterality |
| Throw object | - | 20 | - |
| Touch | Touch Long Other | 470 | Body_part_contact, Body_part_signaller, Laterality, Repetition, |

*Table S2: All significant 2-grams (one antecedent, one consequent) and 3-grams (two antecedents, one consequent) in the 5 second time-window.*

| **Antecedent** | **Consequent** | **Count** | **Conditional Probability** | **Expected Conditional Probability** | **P-Value** |
| --- | --- | --- | --- | --- | --- |
| BigLoudScratch, BigLoudScratch | BigLoudScratch | 21 | 0.553 | 0 | 0 |
| HitObject, StompObject | StompObject | 9 | 0.29 | 0 | 0 |
| ObjectShake, ObjectShake | HitObject | 8 | 0.145 | 0 | 0 |
| HitObject, HitObject | HitObject | 20 | 0.488 | 0 | 0 |
| HitObject, StompObject | HitObject | 11 | 0.355 | 0 | 0 |
| ObjectShake, ObjectShake | ObjectMove | 5 | 0.091 | 0 | 0 |
| ObjectShake, ObjectShake | ObjectShake | 23 | 0.418 | 0 | 0 |
| ObjectShake, StompObject | ObjectShake | 6 | 0.353 | 0 | 0 |
| StompObject, StompObject | ObjectShake | 5 | 0.132 | 0 | 0 |
| StompObject, StompObject | StompObject | 13 | 0.342 | 0 | 0 |
| ObjectMove, ObjectMove | ObjectMove | 5 | 0.263 | 0 | 0 |
| HitObject, ObjectShake | ObjectShake | 8 | 0.444 | 0 | 0 |
| ObjectMove, ObjectShake | ObjectShake | 5 | 0.417 | 0 | 0 |
| StompObject, StompObject | HitObject | 9 | 0.237 | 0 | 0 |
| StompObject, HitObject | HitObject | 11 | 0.367 | 0 | 0 |
| ObjectShake, HitObject | ObjectShake | 6 | 0.3 | 0 | 0 |
| ObjectShake, ObjectShake | Jump | 6 | 0.109 | 0 | 0 |
| HitObject, HitObject | ObjectShake | 5 | 0.122 | 0 | 0 |
| HitObject, ObjectShake | HitObject | 5 | 0.278 | 0 | 0 |
| Push, Push | Push | 16 | 0.762 | 0 | 0 |
| StompObject, HitObject | StompObject | 11 | 0.367 | 0 | 0 |
| Fling, Fling | Fling | 7 | 1 | 0 | 0 |
| HitOther, HitOther | HitOther | 12 | 0.571 | 0 | 0 |
| Touch, Touch | Touch | 13 | 0.565 | 0 | 0 |
| HitObject, HitObject | StompObject | 7 | 0.171 | 0 | 0 |
| Reach, Reach | Reach | 26 | 0.743 | 0 | 0 |
| Pull, Pull | Pull | 24 | 0.857 | 0 | 0 |
| Grab, Grab | Grab | 5 | 0.556 | 0 | 0 |
| Present, BigLoudScratch | BigLoudScratch | 6 | 0.667 | 0 | 0 |
| BigLoudScratch, Present | BigLoudScratch | 16 | 0.889 | 0 | 0 |
| BigLoudScratch, BigLoudScratch | Present | 9 | 0.237 | 0 | 0 |
| BigLoudScratch | BigLoudScratch | 122 | 0.455 | 0 | 0 |
| Present | BigLoudScratch | 107 | 0.856 | 0 | 0 |
| ObjectMove | ObjectMove | 33 | 0.25 | 0 | 0 |
| Stroke | Stroke | 7 | 0.5 | 0 | 0 |
| Embrace | Bite | 16 | 0.64 | 0 | 0 |
| Push | Push | 49 | 0.538 | 0 | 0 |
| HitObject | StompObject | 52 | 0.22 | 0 | 0 |
| StompObject | StompObject | 70 | 0.299 | 0 | 0 |
| ObjectMove | StompObject | 16 | 0.121 | 0 | 0 |
| ObjectShake | ObjectShake | 122 | 0.469 | 0 | 0 |
| ObjectShake | HitObject | 29 | 0.112 | 0 | 0 |
| HitObject | ObjectMove | 11 | 0.047 | 0 | 0 |
| Reach | Reach | 89 | 0.636 | 0 | 0 |
| Reach | Touch | 16 | 0.114 | 0 | 0 |
| Swing | HitObject | 9 | 0.077 | 0 | 0 |
| HitObject | HitObject | 77 | 0.326 | 0 | 0 |
| BigLoudScratch | Present | 84 | 0.313 | 0 | 0 |
| Touch | Touch | 63 | 0.46 | 0 | 0 |
| StompObject | HitObject | 64 | 0.274 | 0 | 0 |
| BigLoudScratch | Pull | 8 | 0.03 | 0 | 0 |
| Shake | Shake | 8 | 0.205 | 0 | 0 |
| Pull | Pull | 40 | 0.606 | 0 | 0 |
| HitObject | Jump | 9 | 0.038 | 0 | 0 |
| Jump | HitObject | 7 | 0.269 | 0 | 0 |
| HitObject | Swing | 9 | 0.038 | 0 | 0 |
| Swing | HitOther | 11 | 0.094 | 0 | 0 |
| ObjectShake | ObjectMove | 20 | 0.077 | 0 | 0 |
| Rocking | ObjectShake | 5 | 0.357 | 0 | 0 |
| ObjectShake | Rocking | 6 | 0.023 | 0 | 0 |
| ObjectMove | ObjectShake | 24 | 0.182 | 0 | 0 |
| ObjectShake | Jump | 8 | 0.031 | 0 | 0 |
| Jump | ObjectShake | 6 | 0.231 | 0 | 0 |
| Push | Touch | 15 | 0.165 | 0 | 0 |
| StompObject | ObjectMove | 14 | 0.06 | 0 | 0 |
| ObjectMove | HitObject | 16 | 0.121 | 0 | 0 |
| Touch | Reach | 7 | 0.051 | 0 | 0 |
| HitObject | HitOther | 9 | 0.038 | 0 | 0 |
| StompObject | Swing | 12 | 0.051 | 0 | 0 |
| Fling | Fling | 21 | 0.677 | 0 | 0 |
| StompObject | ObjectShake | 19 | 0.081 | 0 | 0 |
| ObjectShake | StompObject | 35 | 0.135 | 0 | 0 |
| Dangle | Swing | 28 | 0.315 | 0 | 0 |
| Bite | HitOther | 6 | 0.171 | 0 | 0 |
| Touch | GrabHold | 7 | 0.051 | 0 | 0 |
| Grab | Bite | 18 | 0.254 | 0 | 0 |
| StompObject | Dangle | 10 | 0.043 | 0 | 0 |
| Swing | Shake | 5 | 0.043 | 0 | 0 |
| GrabHold | Bite | 18 | 0.419 | 0 | 0 |
| Swing | Dangle | 13 | 0.111 | 0 | 0 |
| Shake | Swing | 5 | 0.128 | 0 | 0 |
| HitOther | Grab | 12 | 0.09 | 0 | 0 |
| Grab | Grab | 21 | 0.296 | 0 | 0 |
| GrabHold | Grab | 7 | 0.163 | 0 | 0 |
| ObjectMove | Swing | 9 | 0.068 | 0 | 0 |
| Swing | Swing | 24 | 0.205 | 0 | 0 |
| Swing | ObjectMove | 9 | 0.077 | 0 | 0 |
| ObjectMove | Shake | 5 | 0.038 | 0 | 0 |
| ObjectMove | ThrowObject | 5 | 0.038 | 0 | 0 |
| HitObject | ObjectShake | 33 | 0.14 | 0 | 0 |
| Swing | ObjectShake | 12 | 0.103 | 0 | 0 |
| Swing | StompObject | 10 | 0.085 | 0 | 0 |
| ObjectShake | Swing | 15 | 0.058 | 0 | 0 |
| StompObject | HitOther | 7 | 0.03 | 0 | 0 |
| StompObject | HeadStand | 5 | 0.021 | 0 | 0 |
| Dangle | HitObject | 13 | 0.146 | 0 | 0 |
| HitObject | Dangle | 5 | 0.021 | 0 | 0 |
| Dangle | StompObject | 5 | 0.056 | 0 | 0 |
| Bite | Bite | 10 | 0.286 | 0 | 0 |
| PresentGenitals | PresentGenitals | 9 | 0.429 | 0 | 0 |
| Touch | Push | 9 | 0.066 | 0 | 0 |
| Grab | HitOther | 7 | 0.099 | 0 | 0 |
| HitOther | HitOther | 52 | 0.391 | 0 | 0 |
| Pull | Push | 7 | 0.106 | 0 | 0 |
| HitObject | Reach | 5 | 0.021 | 0 | 0 |
| Raise | Touch | 6 | 0.09 | 0 | 0 |
| Reach | Raise | 6 | 0.043 | 0 | 0 |
| Raise | Reach | 6 | 0.09 | 0 | 0 |
| Raise | Raise | 10 | 0.149 | 0 | 0 |
| Dangle | Dangle | 6 | 0.067 | 0 | 0 |
| Grab | Touch | 7 | 0.099 | 0 | 0 |
| Touch | Bite | 8 | 0.058 | 0 | 0 |
| Touch | HitOther | 5 | 0.036 | 0 | 0 |
| LeafClip | LeafClip | 10 | 0.588 | 0 | 0 |
| Dangle | HitOther | 13 | 0.146 | 0 | 0 |
| HitOther | Swing | 8 | 0.06 | 0 | 0 |
| Grab | GrabHold | 6 | 0.085 | 0 | 0 |
| HitOther | Dangle | 5 | 0.038 | 0 | 0 |
| Dangle | Grab | 5 | 0.056 | 0 | 0 |
| LeafClip | ObjectShake | 5 | 0.294 | 0 | 0 |
| HitOther | GrabHold | 5 | 0.038 | 0 | 0 |
| HitOther | StompOther | 8 | 0.06 | 0 | 0 |
| Touch | Grab | 5 | 0.036 | 0 | 0 |
| Raise | StompObject | 5 | 0.075 | 0 | 0 |
| Pull | Touch | 5 | 0.076 | 0 | 0 |
| Raise | BigLoudScratch | 23 | 0.343 | 0 | 0 |
| HitObject | BigLoudScratch | 6 | 0.025 | 0 | 0 |
| Touch | Raise | 7 | 0.051 | 0 | 0 |
| BigLoudScratch | Raise | 14 | 0.052 | 0 | 0 |
| BigLoudScratch | Push | 14 | 0.052 | 0 | 0 |
| BigLoudScratch | PresentGenitals | 5 | 0.019 | 0 | 0 |
| Push | Raise | 6 | 0.066 | 0 | 0 |

*Table S3: All significant 2-grams (one antecedent, one consequent) and 3-grams (two antecedents, one consequent) in the rapid fire time-window.*

| **Antecedent** | **Consequent** | **Count** | **Conditional Probability** | **Expected Conditional Probability** | **P-Value** |
| --- | --- | --- | --- | --- | --- |
| HitObject, StompObject | HitObject | 8 | 0.533 | 0 | 0 |
| ObjectShake, ObjectShake | ObjectShake | 7 | 0.5 | 0 | 0 |
| StompObject, StompObject | StompObject | 7 | 0.35 | 0 | 0 |
| HitObject, HitObject | HitObject | 9 | 0.45 | 0 | 0 |
| StompObject, StompObject | HitObject | 6 | 0.3 | 0 | 0 |
| StompObject, HitObject | HitObject | 9 | 0.429 | 0 | 0 |
| StompObject, HitObject | StompObject | 10 | 0.476 | 0 | 0 |
| HitOther, HitOther | HitOther | 8 | 0.571 | 0 | 0 |
| HitObject, HitObject | StompObject | 6 | 0.3 | 0 | 0 |
| Pull, Pull | Pull | 11 | 0.917 | 0 | 0 |
| BigLoudScratch, Present | BigLoudScratch | 9 | 1 | 0 | 0 |
| Present | BigLoudScratch | 93 | 0.903 | 0 | 0 |
| ObjectMove | ObjectMove | 19 | 0.253 | 0 | 0 |
| StompObject | StompObject | 48 | 0.291 | 0 | 0 |
| ObjectMove | StompObject | 15 | 0.2 | 0 | 0 |
| ObjectShake | HitObject | 18 | 0.138 | 0 | 0 |
| Swing | HitObject | 6 | 0.088 | 0 | 0 |
| BigLoudScratch | Present | 52 | 0.486 | 0 | 0 |
| Touch | Touch | 33 | 0.55 | 0 | 0 |
| Reach | Touch | 6 | 0.24 | 0 | 0 |
| StompObject | HitObject | 56 | 0.339 | 0 | 0 |
| HitObject | StompObject | 41 | 0.275 | 0 | 0 |
| Jump | HitObject | 7 | 0.28 | 0 | 0 |
| Swing | HitOther | 6 | 0.088 | 0 | 0 |
| Stroke | Stroke | 5 | 0.625 | 0 | 0 |
| HitObject | HitObject | 44 | 0.295 | 0 | 0 |
| ObjectShake | ObjectShake | 46 | 0.354 | 0 | 0 |
| ObjectShake | ObjectMove | 9 | 0.069 | 0 | 0 |
| ObjectMove | ObjectShake | 10 | 0.133 | 0 | 0 |
| Jump | ObjectShake | 5 | 0.2 | 0 | 0 |
| Push | Touch | 5 | 0.143 | 0 | 0 |
| StompObject | ObjectMove | 7 | 0.042 | 0 | 0 |
| ObjectMove | HitObject | 11 | 0.147 | 0 | 0 |
| HitObject | ObjectMove | 7 | 0.047 | 0 | 0 |
| Fling | Fling | 6 | 0.667 | 0 | 0 |
| StompObject | ObjectShake | 13 | 0.079 | 0 | 0 |
| Dangle | Swing | 21 | 0.396 | 0 | 0 |
| BigLoudScratch | BigLoudScratch | 34 | 0.318 | 0 | 0 |
| ObjectShake | StompObject | 26 | 0.2 | 0 | 0 |
| Grab | Bite | 18 | 0.5 | 0 | 0 |
| GrabHold | Bite | 16 | 0.615 | 0 | 0 |
| Swing | Dangle | 10 | 0.147 | 0 | 0 |
| Pull | Pull | 24 | 0.667 | 0 | 0 |
| HitOther | Grab | 9 | 0.1 | 0 | 0 |
| Grab | Grab | 8 | 0.222 | 0 | 0 |
| HitObject | ObjectShake | 26 | 0.174 | 0 | 0 |
| ObjectShake | Swing | 11 | 0.085 | 0 | 0 |
| StompObject | HitOther | 5 | 0.03 | 0 | 0 |
| Dangle | HitObject | 8 | 0.151 | 0 | 0 |
| ObjectShake | Jump | 7 | 0.054 | 0 | 0 |
| Swing | ObjectShake | 7 | 0.103 | 0 | 0 |
| HitObject | Jump | 7 | 0.047 | 0 | 0 |
| Push | Push | 15 | 0.429 | 0 | 0 |
| Swing | Swing | 11 | 0.162 | 0 | 0 |
| Swing | StompObject | 7 | 0.103 | 0 | 0 |
| Pull | Push | 5 | 0.139 | 0 | 0 |
| HitOther | HitOther | 36 | 0.4 | 0 | 0 |
| Raise | Raise | 7 | 0.156 | 0 | 0 |
| Reach | Reach | 10 | 0.4 | 0 | 0 |
| StompObject | Swing | 7 | 0.042 | 0 | 0 |
| Touch | HitOther | 5 | 0.083 | 0 | 0 |
| Dangle | HitOther | 9 | 0.17 | 0 | 0 |
| StompObject | Dangle | 7 | 0.042 | 0 | 0 |
| HitOther | StompOther | 8 | 0.089 | 0 | 0 |
| Raise | BigLoudScratch | 19 | 0.422 | 0 | 0 |
| Embrace | Bite | 12 | 0.75 | 0 | 0 |
| BigLoudScratch | Push | 7 | 0.065 | 0 | 0 |
| BigLoudScratch | Raise | 7 | 0.065 | 0 | 0 |

*Table S4: All significant 2-grams (one antecedent, one consequent) in the overlap time-window.*

| **Antecedent** | **Consequent** | **Count** | **Conditional Probability** | **Expected Conditional Probability** | **P-Value** |
| --- | --- | --- | --- | --- | --- |
| Present | BigLoudScratch | 89 | 0.937 | 0 | 0 |
| StompObject | StompObject | 27 | 0.255 | 0 | 0 |
| ObjectShake | HitObject | 18 | 0.194 | 0 | 0 |
| ObjectMove | ObjectMove | 5 | 0.161 | 0 | 0 |
| BigLoudScratch | Present | 39 | 0.619 | 0 | 0 |
| StompObject | HitObject | 43 | 0.406 | 0 | 0 |
| HitObject | StompObject | 28 | 0.318 | 0 | 0 |
| Jump | HitObject | 7 | 0.304 | 0 | 0 |
| HitObject | HitObject | 23 | 0.261 | 0 | 0 |
| ObjectShake | ObjectShake | 28 | 0.301 | 0 | 0 |
| ObjectShake | ObjectMove | 5 | 0.054 | 0 | 0 |
| Jump | ObjectShake | 5 | 0.217 | 0 | 0 |
| StompObject | ObjectMove | 5 | 0.047 | 0 | 0 |
| ObjectMove | HitObject | 5 | 0.161 | 0 | 0 |
| StompObject | ObjectShake | 7 | 0.066 | 0 | 0 |
| Dangle | Swing | 15 | 0.429 | 0 | 0 |
| BigLoudScratch | BigLoudScratch | 11 | 0.175 | 0 | 0 |
| ObjectShake | StompObject | 22 | 0.237 | 0 | 0 |
| Grab | Bite | 18 | 0.9 | 0 | 0 |
| GrabHold | Bite | 15 | 0.789 | 0 | 0 |
| HitObject | ObjectShake | 17 | 0.193 | 0 | 0 |
| ObjectShake | Swing | 8 | 0.086 | 0 | 0 |
| ObjectMove | StompObject | 11 | 0.355 | 0 | 0 |
| HitObject | Jump | 6 | 0.068 | 0 | 0 |
| Swing | StompObject | 7 | 0.163 | 0 | 0 |
| HitOther | HitOther | 13 | 0.342 | 0 | 0 |
| Touch | Touch | 6 | 0.545 | 0 | 0 |
| Swing | Dangle | 7 | 0.163 | 0 | 0 |
| Dangle | HitOther | 7 | 0.2 | 0 | 0 |
| Swing | Swing | 5 | 0.116 | 0 | 0 |
| HitOther | StompOther | 5 | 0.132 | 0 | 0 |
| Embrace | Bite | 11 | 1 | 0 | 0 |
| Raise | BigLoudScratch | 13 | 0.542 | 0 | 0 |
| BigLoudScratch | Push | 5 | 0.079 | 0 | 0 |
| BigLoudScratch | Raise | 6 | 0.095 | 0 | 0 |

*Table S5: All significant 2-grams (one antecedent, one consequent) in the overlap time-window.*

| **Antecedent** | **Consequent** | **Count** | **Conditional Probability** | **Expected Conditional Probability** | **P-Value** |
| --- | --- | --- | --- | --- | --- |
| BigLoudScratch, BigLoudScratch | BigLoudScratch | 8 | 0.615 | 0 | 0 |
| HitObject, HitObject | HitObject | 6 | 0.5 | 0 | 0 |
| ObjectShake, ObjectShake | ObjectShake | 6 | 0.5 | 0 | 0 |
| Fling, Fling | Fling | 5 | 1 | 0 | 0 |
| Reach, Reach | Reach | 20 | 0.769 | 0 | 0 |
| Push, Push | Push | 5 | 0.556 | 0 | 0 |
| BigLoudScratch | BigLoudScratch | 66 | 0.545 | 0 | 0 |
| Reach | Reach | 70 | 0.707 | 0 | 0 |
| Reach | Touch | 8 | 0.081 | 0 | 0 |
| HitObject | HitObject | 24 | 0.436 | 0 | 0 |
| Push | Push | 22 | 0.579 | 0 | 0 |
| BigLoudScratch | Pull | 5 | 0.041 | 0 | 0 |
| ObjectShake | ObjectShake | 50 | 0.633 | 0 | 0 |
| ObjectShake | ObjectMove | 7 | 0.089 | 0 | 0 |
| StompObject | StompObject | 6 | 0.194 | 0 | 0 |
| ObjectMove | ObjectMove | 9 | 0.22 | 0 | 0 |
| Touch | Touch | 23 | 0.404 | 0 | 0 |
| GrabHold | Grab | 5 | 0.385 | 0 | 0 |
| ObjectMove | ObjectShake | 11 | 0.268 | 0 | 0 |
| ObjectMove | Swing | 5 | 0.122 | 0 | 0 |
| StompObject | HitObject | 6 | 0.194 | 0 | 0 |
| PresentGenitals | PresentGenitals | 6 | 0.545 | 0 | 0 |
| Swing | Swing | 7 | 0.28 | 0 | 0 |
| HitOther | HitOther | 10 | 0.5 | 0 | 0 |
| Fling | Fling | 10 | 0.667 | 0 | 0 |
| Present | BigLoudScratch | 11 | 0.688 | 0 | 0 |
| Grab | Grab | 9 | 0.36 | 0 | 0 |
| Grab | Touch | 5 | 0.2 | 0 | 0 |
| LeafClip | LeafClip | 8 | 0.615 | 0 | 0 |
| Dangle | Swing | 6 | 0.261 | 0 | 0 |
| BigLoudScratch | Present | 21 | 0.174 | 0 | 0 |
| HitObject | BigLoudScratch | 6 | 0.109 | 0 | 0 |
| BigLoudScratch | Raise | 5 | 0.041 | 0 | 0 |
| BigLoudScratch | Push | 7 | 0.058 | 0 | 0 |
| Push | Touch | 5 | 0.132 | 0 | 0 |
